# Opposing spatial gradients of inhibition and neural activity in mouse olfactory cortex

**DOI:** 10.1101/152975

**Authors:** Adam M. Large, Nathan W. Vogler, Martha Canto-Bustos, Paul Schick, Anne-Marie M. Oswald

## Abstract

The spatial representation of stimuli in primary sensory cortices is a convenient scaffold for elucidating the circuit mechanisms underlying sensory processing. In contrast, the anterior piriform cortex (APC) lacks topology for odor identity and appears homogenous in terms of afferent and intracortical excitatory circuitry. Here, we show that an increasing rostral-caudal (RC) gradient of inhibition onto pyramidal cells is commensurate with a decrease in active neurons along the RC axis following exploration of a novel odor environment. This inhibitory gradient is supported by somatostatin interneurons that provide an opposing, rostrally-biased, gradient of inhibition to interneurons. Optogenetic or chemogenetic modulation of somatostatin cells neutralizes the inhibitory gradient onto pyramidal cells. This suggests a novel circuit mechanism whereby opposing spatial gradients of inhibition and disinhibition regulate neural activity along the RC-axis. These findings challenge our current understanding of the spatial profiles of neural circuits and odor processing within APC.

---

It is well established that the spatial organization of sensory information plays an important role in neocortical sensory processing. The retinotopic, tonotopic and somatotopic maps established at the periphery form the basis of stimulus representation in primary visual, auditory and somatosensory cortices. This spatial organization is perhaps the oldest and best understood feature of sensory codes.

In the olfactory system, odor components are encoded by individual olfactory receptor neurons (ORNs) that express a single receptor gene. All ORNs expressing the same receptor project axons to ~2 target glomeruli in the olfactory bulb (OB) ^1,2^. Within the OB, individual mitral/tufted (M/T) cells extend apical dendrites to a single glomerulus ^3^ and respond selectively to glomerular activation ^4,5^. This extreme connection specificity produces a discrete spatial organization of odor information within the OB ^6-9^. However, just one synapse away in the anterior piriform cortex (APC), any semblance of spatial representation for odor identity is lost.

The piriform cortex is a trilaminar cortex that extends along the rostral-caudal (RC) axis of the ventral rodent brain. The two main subdivisions, anterior (APC) and posterior (PPC) piriform cortex, differ with respect to afferent and efferent projections ^10-12^ as well as functional roles in olfactory processing ^13-16^. However, despite the fact that each region comprises ~1-2 mm of the RC axis, odor processing within APC or PPC is considered spatially homogenous. The APC is delineated by the lateral olfactory tract (LOT) that delivers odor information directly from the OB. Single M/T cell axons branch extensively along the LOT ^17-19^ resulting in diffuse pattern of afferent excitation. Likewise, recurrent connections between principal neurons within APC extend over millimeter distances ^20,21^. Consistent with this distributed excitatory architecture, there is no topography for odor identity in APC. Neurons responsive to a single odor are distributed along the RC-axis of the APC ^22-24^ and nearby neurons respond to different odors ^24-27^. The absence of an “odortopic” map, suggests that, unlike sensory neocortex, space is not a dimension for odor coding in APC.

Nonetheless, there is evidence that odor evoked responses vary along the rostral-caudal axis of APC. For example, odor evoked activity at rostral sites is denser ^22^, has lower concentration thresholds ^28^, and has earlier response times ^24,29-31^ versus caudal sites. Further, in contrast to excitation, intracortical inhibition is asymmetric along the RC axis ^32^. Finally, rostral APC projects more densely to the OB and orbitofrontal cortex (OFC) while caudal APC and PPC project to agranular insula (AI) ^11,33-35^. Thus, contrary to preconceived notions, space may be a relevant feature of olfactory processing in APC. However, a central challenge is elucidating the circuit mechanisms and functional roles of RC spatial patterning in APC.

In this study, we use optogenetic ^36^ and chemogenetic tools ^37^ as well as targeted-recombination in active populations (TRAP) ^38^ to investigate the circuitry underlying the spatial profiles of inhibition and neural activity in APC. Specifically, we find that inhibition of pyramidal cells (PC) increases on a millimeter scale from rostral to caudal APC. Our findings suggest a disinhibitory circuit mechanism, mediated by somatostatin interneurons, underlies the RC patterning of inhibition. This increasing gradient of inhibition is commensurate with a context dependent, RC decrease in the density of active neurons during odor exposure. Moreover we show that spatial patterning of inhibition and neural activity is laminar dependent. Altogether, our findings provide new insights to the RC spatial patterns of inhibition and neural activity in APC and suggest space is a dimension of olfactory processing in cortex.

## Results

### Asymmetric inhibitory circuitry in APC

Spatially asymmetric inhibition of pyramidal cells (PC) in APC ^32^ has been previously demonstrated using glutamate uncaging. Specifically, light evoked uncaging at caudal stimulation sites yielded stronger inhibition of PCs compared to rostral sites. These findings are seemingly at odds with the spatial profiles of excitation ^17-21^ and odor-evoked responses ^26,27^ that do not vary with space. This led us to question, “How and why do inhibitory spatial asymmetries exist in APC?”

Since previous un-caging methods could activate both excitatory and inhibitory neurons, we investigated whether inhibitory circuits alone are sufficient to reproduce inhibitory asymmetries. Whole cell recordings were made from L2 principal excitatory neurons, namely semilunar cells (SL) and superficial PCs (sPC) as well as deep PCs (dPC) in L3. Recorded neurons were centrally located along the RC axis in sagittal slices of APC. Interneurons were specifically activated using Channelrhodopsin ChR2) expressed under the promoter for vesicular GABA transporter (VGAT). VGAT-ChR2 interneurons were stimulated using restricted spots (~70 µm diameter) of blue light in 4x5 grid surrounding the recorded cell (schematic, **Figure 1A, D1** ^39^). The inhibitory strength was quantified as the area (pAs) under IPSC evoked at each stimulation site **(Figure 1B1,C1, D2)**. We have previously reported the inhibitory strength by cell type and layer ^39^. Here, we quantify the asymmetry in inhibitory strength evoked at rostral versus caudal sites. An asymmetric bias index was calculated as the difference in average inhibition at caudal versus rostral stimulation sites (I_C_-I_R_), divided by the sum of the inhibition from both sides (I_C_+I_R_), (**Figure 1B2,C2,D3**). Thus, solely caudal inhibition produces a bias value of +1 while -1 corresponds to rostral inhibition. Inhibition was not significantly asymmetric in SL cells (bias: 0.08 ± 0.11, p: 0.49, n=13, **Figure 1B2**). However, sPCs (L2) and dPCs (L3) received significant, caudally biased inhibition (sPC bias: 0.22 ± 0.08, p: 0.016, n=14, **Figure 1C2**; dPC bias: 0.19 ± 0.05; p: 0.0012, n=16, one sample t-test, **Figure 1D3**). Thus, inhibitory circuitry alone is sufficient to reproduce asymmetric inhibition of PCs along the RC-axis. Since we have previously found that inhibition of sPCs is significantly weaker than dPCs, and inhibition from L3 is stronger than L2 ^39^, we focused the remainder of our experiments on L3 dPCs.

**Figure 1.**
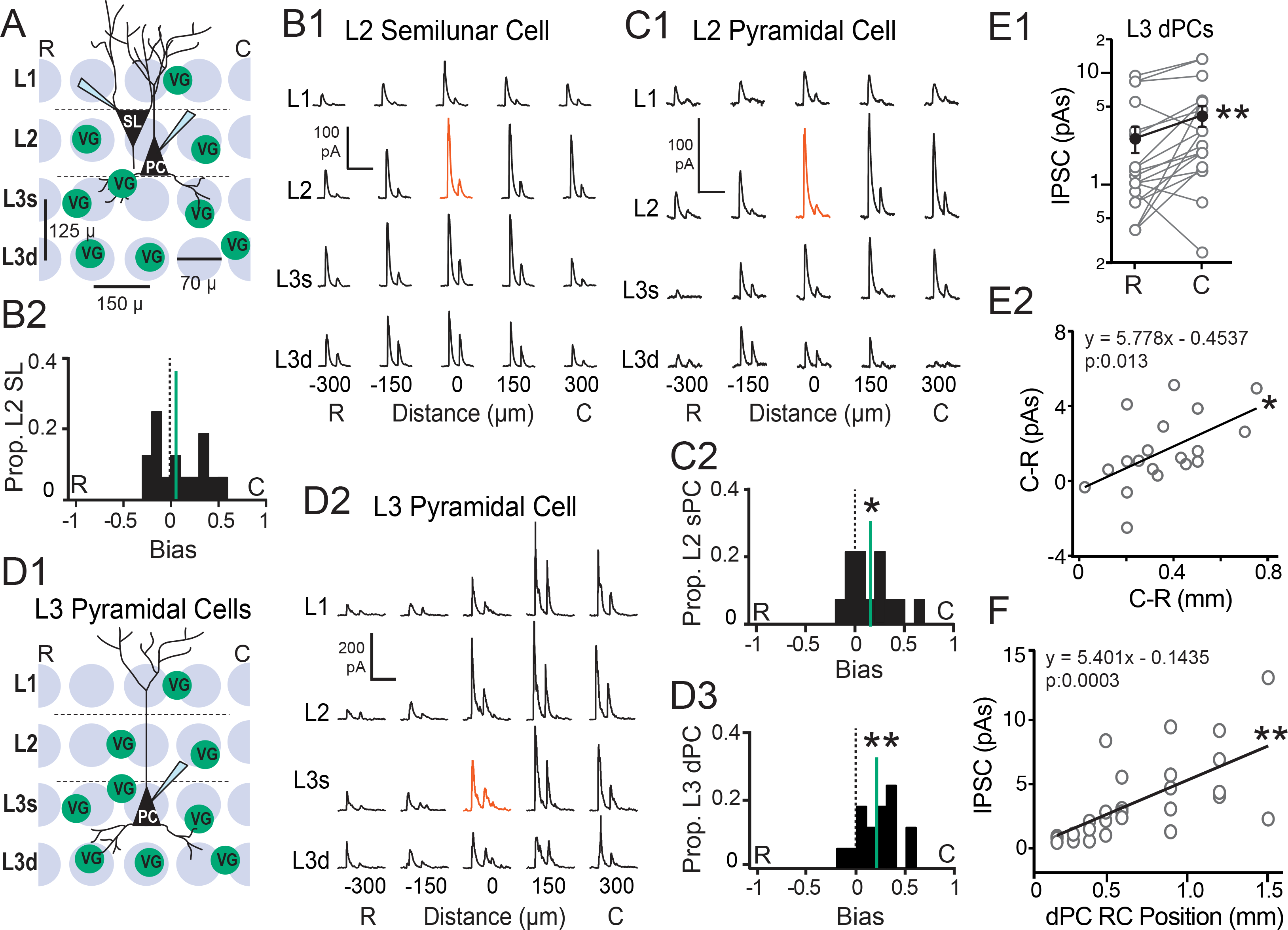
Caudally biased, asymmetric inhibition of pyramidal cells in APC. **A)** Schematic of grid-stimulation paradigm for L2 excitatory neurons-semilunar cells (SL) and superficial (s) pyramidal cells (sPCs). VG-interneuron expressing ChR2 under VGAT promoter. R: rostral, C: caudal. **B1)** IPSCs recorded during focal light stimulation at each grid location in a representative SL cell. Scale bars: vertical 100 pA, horizontal 200 ms. Red trace indicates location of recorded cell. **B2)** Bias values are uniformly distributed across SL cells. Negative values correspond to greater average inhibition from rostral sites and positive values correspond to greater inhibition from caudal sites. **C1)** IPSCs recorded from a representative L2 sPC. **C2)** Predominantly positive bias values in sPCs indicates stronger inhibition from caudal versus rostral sites (* p<0.05, n=14, one sample t-test). Scale bars: vertical 100 pA, horizontal 200 ms. **D1)** Schematic of grid-stimulation paradigm for L3 deep pyramidal cells (dPCs). **D2)** IPSCs recorded from a representative L3 dPC. Scale bars: vertical 200 pA, horizontal 200 ms. **D3)** Predominantly positive bias values in dPCs indicates stronger inhibition from caudal versus rostral sites (** p<0.01, n=16, one sample t-test). **E1)** Recordings of two dPCs in the same slice indicate that the caudal (C) neuron of a pair receives stronger inhibition than the rostral (R) neuron (**p<0.01, n=19 pairs, paired t-test). IPSCs were evoked using a 70 µm light spot centered on the soma of the recorded PC. **E2)** The difference in inhibition (pAs) between the caudal (C) and rostral (R) cell in each pair (E1) is plotted against the difference in RC distance between the two cells. As the distance between the two cells increases, the difference in inhibition also increases (* p<0.05, linear regression, n=19). **F)** Inhibitory strength versus RC position of the dPC relative to the rostral start of the LOT in the sagittal slice. Inhibition increases with RC distance (**p<0.01, linear regression, n=27).

One interpretation of these findings is that caudally located dPCs receive stronger inhibition than rostral dPCs. To investigate this possibility, we compared local inhibitory strength in dPC pairs (n=19) separated by (100-1000 µm) distances along the ~1.5 mm RC-axis. On average, the caudal neuron of the pair received significantly stronger inhibition (4.18 ± 0.90 pAs) than the rostral neuron (2.60 ± 0.68 pAs, p: 0.002, paired t-test, **Figure 1E1**). Moreover, as the distance between the rostral and caudal neuron increased, the difference in inhibitory strength increased (slope: 5.8 ± 2.1 pAs/mm, p: 0.013, R^2^: 0.32, **Figure 1E2**). Finally, across the population of dPCs, inhibitory strength increased with soma location relative to the rostral start of the LOT in the sagittal slice (slope: 5.4 ± 1.3 pAs/mm, p: 0.0003, n=27, R^2^: 0.41, **Figure 1F**). These findings demonstrate an increasing gradient of inhibition onto dPCs with a spatial scale that extends >1.0 mm of the RC axis of APC.

To test if stronger caudal inhibition is a general feature of APC we investigated inhibition of L2 and L3 interneurons using grid stimulation of VGAT-ChR2 interneurons (**Figure 2A**). L2 interneurons received weaker inhibition (3.42 ± 0.5 pA) than L3 interneurons (7.02 ± 0.92 pAs, p: 0.0043, ANOVA-Tukey, **Figure 2B**). In addition, L2 interneurons did not receive significantly asymmetric inhibition (bias: 0.04 ± 0.07, p: 0.11, n=18, **Figure 2C1, 2**). L3 interneurons received comparable inhibition to dPCs (8.63 ± 0.84, p: 0.239, ANOVA-Tukey, **Figure 2B**). However, L3 interneurons received stronger inhibition from rostral rather than caudal sites (bias: -0.13 ± 0.05, p: 0.013, n=25, **Figure 2D1,2**) and bias indices of L3 interneurons significantly differed from dPCs (p: 0.0001, t-test, **Figure 2E**). Thus, dPCs and L3 interneurons have opposing spatial profiles of inhibition-dPCs receive stronger inhibition from caudal sites versus rostral sites (I_C_: 0.63 ± 0.034, I_R_: 0.44 ± 0.042 p: 0.003, paired t-test) and L3 interneurons receive significantly more inhibition from rostral sites versus caudal (I_R_: 0.54 ± 0.033, I_C:_ 0.42 ± 0.034 p: 0.008, paired t-test **Figure 2F**).

**Figure 2.**
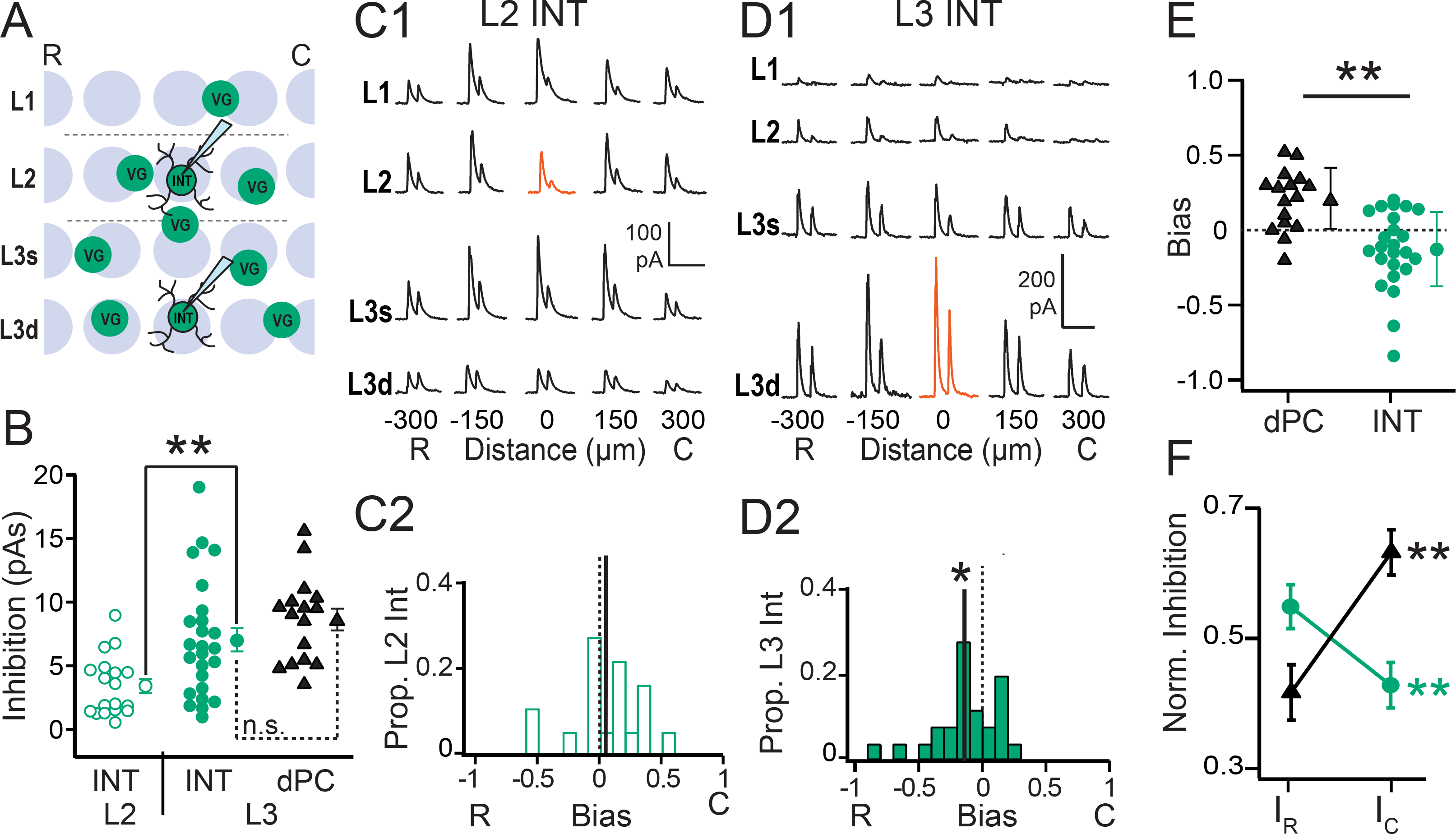
Rostrally biased, asymmetric inhibition of L3 interneurons. **A)** Schematic of grid-stimulation paradigm for L2 and L3 inhibitory neurons (INT). **B)** Inhibition is significantly stronger in L3 INT (filled green circles) than L2 INT (** p<0.01, ANOVA, open green circles) but did not significantly differ from L3 dPCs (solid black triangles). Values correspond to the maximum IPSC for each cell regardless of location in grid. **C1)** IPSCs recorded during focal light stimulation at each grid location in a representative L2 INT. Scale bars: vertical 100 pA, horizontal 200 ms. Red trace indicates location of recorded cell. **C2)** The bias values for L2 INTs do not differ from zero. **D1)** IPSCs recorded from a representative L3 INT. Scale bars: vertical 200 pA, horizontal 200 ms. **D2)** Asymmetric distribution of bias values in dPCs indicates stronger inhibition from rostral versus caudal sites (** p<0.05, n=25, one sample t-test). **E)** The bias values of dPCs (black triangles) significantly differed from L3 INTs (** p<0.01, unpaired t-test, green circles). **F)** Average inhibitory strength across rostral (I_R_) or caudal (I_C_) sites in dPCs (black triangles) and L3 INT (green circles). Opposing inhibitory asymmetries: in dPCs, average I_C_ was significantly greater than I_R_ (black ** p<0.01 paired t-test), while I_R_ was significantly greater than I_C_ l in L3 INTs (green ** p<0.01, paired t-test).

How do inhibitory circuits implement opposing RC inhibitory asymmetries and what is the functional role of this opposition? A simple mechanism to increase inhibition is to increase the number of interneurons. Since the majority (~85%) of PC-targeting interneurons in L3 express parvalbumin (PV), somatostatin (SST) and/or calbindin (CB) ^40^, we investigated the RC distributions of these three interneuron classes. SST cells were identified as tdTom(+) cells in SST-Ai14 mice (**Figure 3A**, n=7 mice) while PV and CB cells were identified using anti-CB (**Figure 3B**, n=6 mice) or anti-PV immunochemistry (**Figure 3C**, n=6 mice). To quantify interneuron density, somas were counted in L3 along 1 to 1.5 mm (0.1 mm increments) of the RC axis (**Figure 3A1-C1**, ROIs dashed areas). As previously shown ^40^, the average densities (cells/mm^2^) of SST (235 ± 14) and CB cells (174 ± 27) were greater than PV cells (109 ± 9.0, p: 0.0054, KW-test). To investigate RC patterning, densities for each section were normalized to the most rostral section (~2.46 mm from Bregma ^41^). For each animal, normalized density versus RC distance was linearly fit to obtain the slope of the change in density per mm^-1^ (**Figure 3A1-C2**, **Table 1**). Only SST-cell densities consistently varied along the RC axis (**Figure 3A1-4, Table 1**). In the majority of mice (n=5/7), linear fits of SST cell density had significantly negative slope values (mean slope: -0.20 ± 0.04, p: 0.002, MWU-test, **Figure 3A4**, **Table 1**), and the average normalized density across all animals (red filled circles, **Figure 3A3**) also decreased (slope: -0.25 ± 0.03, p<0.0001). The RC distributions of CB cells were highly variable with individual mice showing increases (n=2, i.e. **Figure 3B1,2**), decreases (n=1) or no change (n=3) along the RC axis. The distribution of CB slopes did not differ from zero (0.13 ± 0.16, p: 0.37, MW-test, **Figure B4**) and the average normalized density did not change with RC distance (slope: 0.11 ± 0.17, p: 0.55, **Figure 3B3**). PV cell density appears to decrease along the RC axis. However, this was only significant in one PV animal (filled black circle, **Figure 3C4, Table 1**) and the average normalized density across all animals (-0.27 ± 0.10, p: 0.031, **Figure 3C3**). In the majority of mice (n=5/6), PV cell density did not significantly vary with RC distance and the distribution of slopes did not differ from zero (-0.16 ± 0.08, p: 0.065, MW-test, open circles, **Figure 3C4**). Altogether, these findings suggest it is unlikely that stronger caudal inhibition of dPCs arises from a rostral-to-caudal increase in interneuron density in L3.

**Figure 3.**
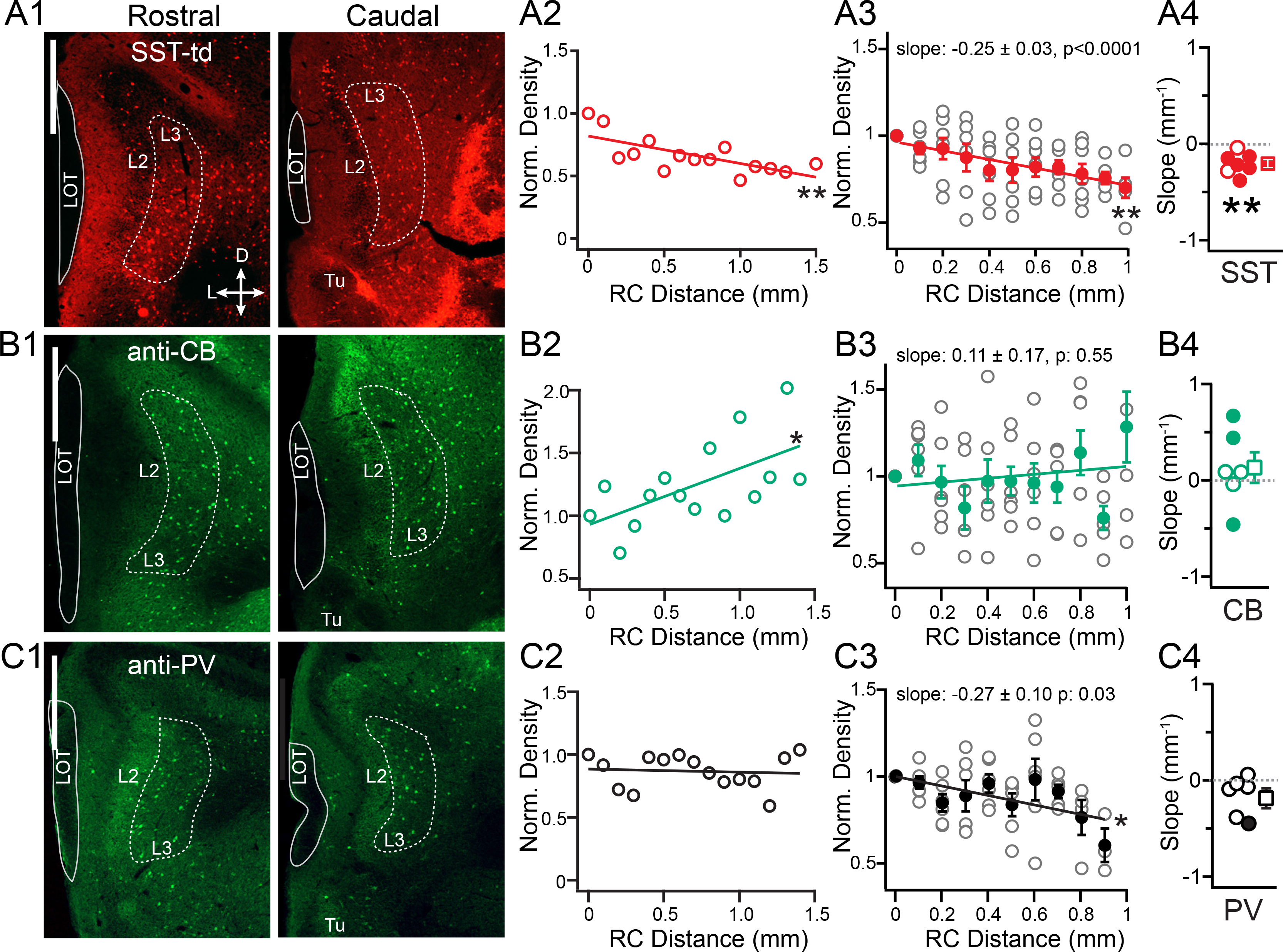
Rostral-caudal distributions of L3 interneurons. **A1)** Representative coronal sections from rostral (0-200 µm, left) and caudal (within last 300 µm, right) APC showing SST-tdTom(+) cells in L3 region of interest (ROI, dashed line). Scale bar: 500 µm **A2)** Normalized density versus RC distance for the mouse shown in A1 (SST-mouse #2, **Table 1**, ** p<0.01. **A3)** Normalized density of SST-cells versus distance for all mice (open circles). The average (± SE, n=7) normalized density of SST cells across mice significantly decreased with RC distance (red circles, **p<0.01). **A4)** Distribution of slopes from linear fits to data from individual mice. Solid circles indicate significantly negative slopes (p<0.05). The distribution of slope values was significantly non-zero (** p: 0.002 MWU-test). **B1-4)** As in A1-4, except for Calbindin(+) interneurons (CB). **B1,2)** Data from CB mouse #5 in **Table 1**. **B3**) On average, there is no change in density of CB cells along the RC axis (filled green circles, p: 0.55). **B4)** In individual mice CB cells significantly increased or decreased (filled green circles) along the RC axis, but the distribution of slopes did not differ from zero (p: 0.37, MWU-test). **C1-4)** As in A1-4, except for Parvalbumin(+) interneurons (PV). **B1,2)** Data from PV mouse #6 in **Table 1**. **C3**) On average, the density of PV cells decreased along the RC axis (filled black circles, p: 0.03). **C4)** However, only one mouse showed a significant decrease in PV cells along the RC axis (filled black circle) and the distribution of slopes did not differ from zero (p: 0.07, MWU-test).

**Table 1.** Linear regression slope values for fits to normalized density of three classes of interneuron versus RC distance in individual mice. Bold values correspond to significant (p<0.05) slope values (mm^-1^). Linear regression was also performed on the average normalized density across mice versus distance for each interneuron class (Fit Average). P-values correspond to tests for slope not equal to zero. Finally, the distribution of slope values was compared to zero using a non-parametric Mann-Whitney test (distribution ≠ 0). SST-somatostatin interneurons, PV-parvalbumin interneurons, CB-Calbindin interneurons.

| Mouse | SST |  |  | PV |  |  | CB |  |  |
| --- | --- | --- | --- | --- | --- | --- | --- | --- | --- |
|  | Slope | R<sup>2</sup> | P | Slope | R<sup>2</sup> | P | Slope | R<sup>2</sup> | P |
| 1 | **-0.13 ± 0.03** | **0.48** | **0.0014** | -0.07 ± 0.20 | 0.02 | 0.7385 | 0.08 ± 0.09 | 0.06 | 0.3827 |
| 2 | **-0.22 ± 0.06** | **0.49** | **0.0038** | -0.09 ± 0.16 | 0.06 | 0.5977 | 0.09 ± 0.21 | 0.01 | 0.6787 |
| 3 | -0.04 ± 0.03 | 0.10 | 0.2092 | -0.38 ± 0.20 | 0.31 | 0.0950 | **0.67 ± 0.21** | **0.47** | **0.0102** |
| 4 | **-0.24 ± 0.03** | **0.85** | **<0.0001** | 0.06 ± 0.20 | 0.01 | 0.7621 | **-0.47 ± 0.07** | **0.79** | **<0.0001** |
| 5 | **-0.15 ± 0.03** | **0.61** | **0.001** | **-0.44 ± 0.14** | **0.56** | **0.0124** | **0.44 ± 0.17** | **0.35** | **0.0212** |
| 6 | **-0.37 ± 0.05** | **0.86** | **<0.0001** | -0.02 ± 0.14 | 0.00 | 0.8698 | -0.04 ± 0.10 | 0.01 | 0.6900 |
| 7 | -0.27 ± 0.17 | 0.25 | 0.1402 |  |  |  |  |  |  |
| Fit Average | **-0.25 ± 0.03** | **0.88** | **<0.0001** | **-0.27 ± 0.10** | **0.46** | **0.0305** | 0.11 ± 0.17 | 0.05 | 0.5538 |
| Slope Dist ≠ 0 | **-0.20 ± 0.04** | - | **0.0021** | **-0.16 ± 0.08** | - | **0.0658** | 0.13 ± 0.16 | - | 0.3700 |

We have previously shown that SST-cells inhibit the majority of L3 interneurons in APC ^42^. This suggests that the high rostral density of SST cells could underlie rostrally-biased inhibition of interneurons. To test this, we selectively expressed ChR2 in SST-cells and repeated grid stimulation while recording IPSCs in L3 interneurons and dPCs (**Figure 4A**). As predicted, SST-cells provided rostrally-biased inhibition to the majority (73%) of L3 interneurons (bias: -0.11 ± 0.04, p: 0.02, n=22, **Figure 4B1,2**). Bias values did not significantly differ between VGAT-ChR2 and SST-ChR2 animals (p: 0.69, unpaired t-test). Thus, activating solely SST-cells replicates the rostrally-biased inhibition (I_R_: 0.58 ± 0.03 > I_C_: 0.048 ± 0.03, p: 0.017, paired t-test, **Figure 4D**) of L3 interneurons seen in VGAT-ChR2 animals. Interestingly, the strength of SST-mediated inhibition onto L3 interneurons also did not significantly differ (6.82 ± 1.4 pAs) from that in VGAT-ChR2 animals (7.02 ± 0.92 pAs, p: 0.91, unpaired t-test). Although one should be cautious comparing between transgenic lines, one interpretation is that SST cells are a major source of rostrally-biased inhibition onto L3 interneurons.

**Figure 4.**
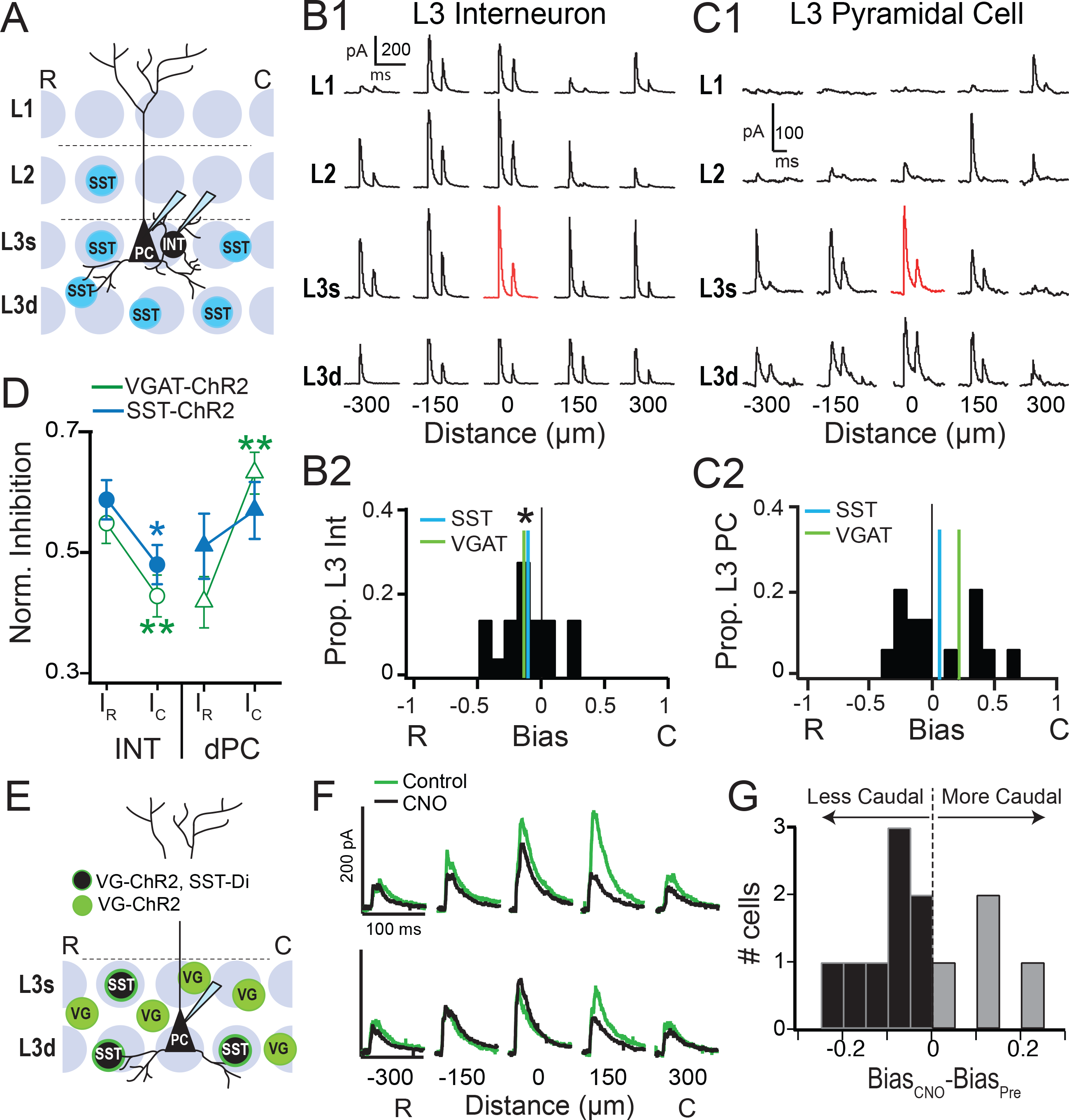
Spatial profiles of SST-cell mediated inhibition of L3 pyramidal cells and interneurons. **A)** Schematic of grid-stimulation paradigm for L3 dPCs and inhibitory neurons (INT) in sagittal sections from SST-ChR2 mice. **B1)** IPSCs recorded during focal light stimulation at each grid location in a representative L3 INT. Scale bars: vertical 200 pA, horizontal 200 ms. Red trace indicates location of recorded cell. **B2)** The distribution of bias values for L3 INTs is significantly rostrally biased (*p<0.05, one sample t-test, blue line) and does not differ from VGAT-ChR2 animals (green line). **C1)** IPSCs recorded from a representative L3 dPC. Scale bars: vertical 100 pA, horizontal 200 ms. **C2)** The distribution of bias values in dPCs is not asymmetric with a mean near zero (blue line) compared to a caudally biased mean in VGAT-ChR2 animals (green line). **D)** Normalized inhibition on rostral versus caudal sides in VGAT-ChR2 (green) and SST-ChR2 (blue) mice is significantly asymmetric and rostrally biased in L3 interneurons (circles, Left). SST-ChR2 mediated inhibition is not significantly asymmetric in dPCs (right triangles, SST-ChR2: blue, VGAT-ChR2: green). **E)** Schematic of optogenetic stimulation of VGAT-ChR2(+) interneurons (green) including SST-cells which also express the inhibitory DREADD, (SST-Di, black with green outline). **F)** IPSCs in response to optogenetic activation of L3 rostral and caudal sites in control (green) and CNO (black). **G)** Change in RC bias (Bias_CNO_-Bias_Pre_) in the presence of CNO. Inhibition became less caudally biased in n=8 cells (negative values, black bars) in CNO but more caudally biased in n=4 cells (positive values, gray bars).

In dPCs, SST-ChR2 mediated inhibition was significantly weaker (4.77 ± 0.89 pAs) than VGAT-ChR2 (8.63 ± 0.84 pAs, p: 0.003, unpaired t-test). This would be expected if SST-cells are only a subset of the interneurons that inhibit dPCs. SST-ChR2 cells provided rostrally-biased inhibition to 50% of dPCs (n=7/14, **Figure 4C1**) compared to only 2/16 in VGAT-ChR2 animals. Consequently, the distribution of bias values was not coherently asymmetric and did not differ from zero (0.06 ± 0.08, p: 0.45, one sample t-test, n=14, **Figure 4C2**). Finally, in contrast to VGAT-ChR2 activation, SST-ChR2 inhibition did not differ between rostral and caudal sites (p: 0.50 paired t-test, **Figure 4D**). Thus, activation of solely SST-cells increases rostrally-biased inhibition of many dPCs and ultimately neutralizes inhibitory bias across the population. These findings are consistent with rostrally-biased distributions of SST-cells and suggest that additional inhibitory circuits are required to produce consistent, caudally biased inhibition of dPCs.

There are two ways to produce caudally biased inhibition of dPCs - 1) increase caudal inhibition; or 2) decrease rostral inhibition. We have not found a mechanism to support increased caudal inhibition. However, we have shown that SST-cells provide rostral inhibition to interneurons (**Figure 4B**), which could decrease rostral inhibition of dPCs. To test this possibility, we bred triple transgenic animals that express ChR2 in all interneurons but only SST-cells express the inhibitory DREADD, hM4Di (abbreviated: VGAT-ChR2-SST-Di). In these mice, the DREADD agonist CNO (20 µM, bath) reduces SST-cell activity. We performed grid stimulation of L3 sites (**Figure 4E**) and compared IPSC strength and RC bias in control conditions (green) versus CNO (black, **Figure 4F**). Upon application of CNO, IPSC strength decreased consistent with a loss of direct, SST mediated inhibition of dPCs (Control: 4.14 ± 0.70 pAs; CNO: 2.72 ± 0.40 pAs, p: 0.007, paired t-test, n=12). If SST-cells influence caudal bias through rostral disinhibition, we expect a loss in SST-mediated inhibition would shift bias to less caudal values. Surprisingly, the mean bias did not change significantly in CNO (bias: +0.17 ± 0.08, n=12) compared to control (bias: +0.19 ± 0.06, 0.57, paired t-test). However, the bias distribution was significantly asymmetric in control conditions (mean>0, p: 0.010, one sample t-test), whereas in CNO, bias values were not significantly asymmetric (p: 0.051, one sample t-test). This is because caudal bias both increased (n=4/12) and decreased (n=8/12) across the dPC population with CNO application. In a small number of dPCs, increased caudal bias (n=4/12, Δ_Bias_= Bias_CNO_-Bias_Control_ =+0.13 ± 0.04, gray bars **Figure 4G**) can be explained by a loss of direct, SST-mediated inhibition at rostral sites. In contrast, the majority of dPCs (n=8/12) shifted toward less caudal bias values with CNO (Δ_Bias_ -0.09 ± 0.03, black bars, **Figure 4G**). In these cells, CNO produced a significantly greater reduction in inhibition at caudal sites (-35 ± 9%) versus rostral sites (-24 ± 7%, p<0.05, WSR-test, n=8). This suggests that rostral interneurons are normally suppressed by SST-cells in control conditions, but rebound during CNO application and neutralize bias. Thus, caudally biased inhibition of dPCs could arise by rostral disinhibition of PCs through SST-to-interneuron microcircuits.

### Rostral-caudal spatial profiles of neural activity

What role might RC patterning of inhibition play in regulating neural activity during olfactory processing? Although odor responsive neurons are distributed in APC, few studies have addressed the spatial patterns of neural activity along the full extent of the APC ^22,24,28^. To investigate the RC patterning of neural activity in APC, we used targeted recombination in active populations (TRAP) in FosCre^ERT^ mice ^38^. These mice express a tamoxifen-dependent cre-recombinase under the promoter for the activity dependent, immediate early gene *c-fos*. We used FosCre^ERT^x Ai14 mice to express the fluorescent protein, tdTomato in neurons activated in the presence hydroxytamoxifen (4-OHT), (**Figure 5A, Supplemental Figure 1**). We measured the spatial distribution of active neurons along the RC axis of APC in three conditions: home cage with no odor (HC), home cage with novel odor (HCO), or exploration of a novel environment with novel odor (NEO) (**Figure 5B1,2**). Briefly, HC (n=6) and HCO (n=6) mice were given a single dose of 4-OHT and then returned to the home cage. HCO mice were allowed to rest for 30 min then exposed to odor in the home cage for 30 min. NEO mice (n=6) were given 4-OHT, rested for 30 min, and then explored a novel environment plus odor for 30 min before being returned to the home cage. The novel environment was a divided arena with two cups of bedding-one odorized, one blank-at the end of each arm (**Figure 5B2**, schematic far right). In a subset of mice (n=4), location within the arena was monitored. NEO mice were highly active and sampled both arms as well as the center (C) of the arena throughout exposure period (30 min, **Figure 5B3**). Mice spent nearly equivalent time per visit in the odorized (8.1 ± 3.5 s) and blank (9.8 ± 3.5 s) arms, but visited the blank arm (49 ± 33 visits) more frequently than the odorized arm (25 ± 14 visits). The majority of HCO and NEO mice were exposed to isoamyl acetate. One cohort of mice (n=2 HCO, n=1 NEO) was exposed to ethyl-butyrate. Results did not differ between odors and were grouped. Following odor exposure, mice remained in their home cages, undisturbed, in the dark for 10-12 hours. Mice were sacrificed 5 days later and tdTom(+) neurons were counted along ~1.5 mm of the RC axis. Neural activity was quantified as the density (cells/mm^2^) of tdTom(+) cells in laminar regions of interest (L2, L3 ROIs, **Figure 5A**) located directly under the LOT. Densities were normalized to the most rostral section for linear fits as described for interneuron densities (**Figure 3**). Summary data is presented in **Figure 5** and **Table 2**; representative mice from each group and odor are shown in **Supplemental Figure 1**.

**Figure 5.**
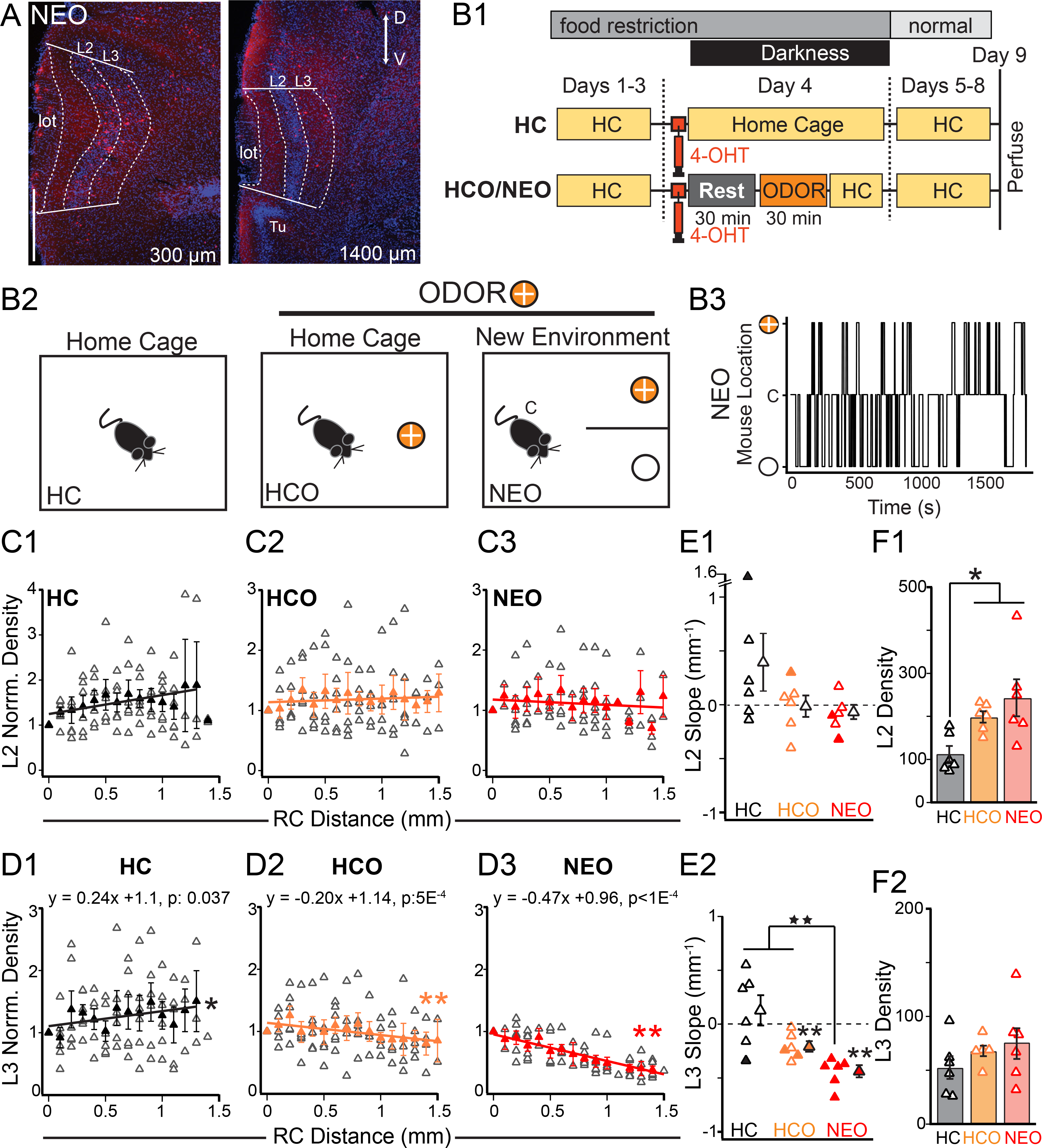
Spatial profiles of neural activity in APC following odor exposure. **A)** tdTom(+) cells from a Fos^ERT^xAi14 mouse in rostral (left) and caudal (right) sections from (NEO condition). Dashed lines delineate Layers 2 and 3 and the lateral olfactory tract (lot). DAPI stain for nuclei (blue). **B1, B2)** Schematics of experimental schedule (**B1**) and contexts **(B2)**. Abbreviations: HC: home cage, HCO: home cage plus odor, NEO: Novel environment plus odor, 4-OHT: 4-hydroxytamoxifen. **B3**) Location of an example mouse in the NEO arena during 30 min odor exposure. Mice continuously move between the odorized (orange circle (+)) and non-odor arms (blank circle (-)) as well as the center (C) of the arena. **C, D)** Normalized density of tdTom(+) cells along the RC axis of L2 **(C)** and L3 (**D).** Open triangles: data from individual animals; filled triangles: average across animals. **E)** Slopes of linear fits to density versus distance for individual mice in each condition in L2 (**E1**) and L3 (**E2**). Filled triangles indicate slopes significantly different from zero (p<0.05, see **Table 1** for p: values). **E2**) In L3, the distribution of slopes in HCO and NEO animals significantly differed from 0 (** p<0.01, MWU-test). Further the distribution of slopes in NEO animals significantly differed from HC and HCO animals (★★ p<0.01, KW-test). **F)** The average density of neurons increases with odor exposure (HCO, NEO) in L2 (**F1, \*** p<0.05, KW-test) but not L3 **(F2)**.

**Table 2.** Linear regression slope values for fits to normalized density of tdTom(+) cells versus RC distance in individual mice. Bold values correspond to significant (p<0.05) slope values (mm^-1^). Underlined values correspond to the cohort of mice exposed to ethyl butyrate (HCO, NEO only) and the remaining mice were exposed isoamyl acetate. Linear regression was also performed on the average normalized density across mice versus distance (Fit Average). P-values correspond to tests for slope not equal to zero. Finally, the distribution of slope values was compared to zero using a non-parametric Mann-Whitney test (distribution ≠ 0). HC-home cage, HCO-home cage plus odor, NEO-novel environment plus odor.

| Layer 2 | HC |  |  | HCO |  |  | NEO |  |  |
| --- | --- | --- | --- | --- | --- | --- | --- | --- | --- |
| Mouse | Slope | R<sup>2</sup> | P | Slope | R<sup>2</sup> | P | Slope | R<sup>2</sup> | P |
| 1 | 0.11 ± 0.21 | 0.03 | 0.6000 | -0.02 ± 0.10 | 0.01 | 0.7864 | **-0.32 ± 0.13** | **0.29** | **0.0276** |
| 2 | -0.14 ± 0.12 | 0.09 | 0.2700 | -0.40 ± 0.37 | 0.09 | 0.3000 | 0.17 ± 0.26 | 0.05 | 0.5100 |
| 3 | 0.60 ± 0.52 | 0.12 | 0.2700 | -0.17 ± 0.11 | 0.18 | 0.1250 | -0.18 ± 0.17 | 0.12 | 0.3220 |
| 4 | **1.62 ± 0.38** | **0.61** | **0.0009** | **0.30 ± 0.13** | **0.27** | **0.0370** | -0.08 ± 0.08 | 0.12 | 0.3297 |
| 5 | 0.22 ± 0.31 | 0.04 | 0.5300 | <u>0.10 ± 0.27</u> | 0.01 | 0.7000 | **-0.10 ± 0.02** | **0.63** | **0.0032** |
| 6 | <u>-0.06 ± 0.13</u> | 0.02 | 0.6400 | <u>0.07 ± 0.10</u> | 0.04 | 0.4800 | <u>0.0008 ± 0.07</u> | 0.00 | 0.9914 |
| Fit Average | **0.42 ± 0.10** | **0.58** | **0.0016** | -0.06 ± 0.06 | 0.08 | 0.2972 | -0.09 ± 0.09 | 0.06 | 0.3500 |
| Slope Dist ≠ 0 | 0.39 ± 0.27 | - | 0.3789 | 0.01 ± 0.10 | - | 0.3789 | -0.08 ± 0.07 | - | 0.0658 |

**Table 2: Linear regression slope values for fits to normalized density of tdTom(+) cells versus RC distance in individual mice.**
| Layer 3 | HC |  |  | HCO |  |  | NEO |  |  |
| --- | --- | --- | --- | --- | --- | --- | --- | --- | --- |
| Mouse | Slope | R<sup>2</sup> | P | Slope | R<sup>2</sup> | P | Slope | R<sup>2</sup> | P |
| 1 | **-0.34 ± 0.12** | **0.42** | **0.0224** | **-0.24 ± 0.10** | **0.32** | **0.0330** | **-0.68 ± 0.08** | **0.84** | **<0.0001** |
| 2 | -0.16 ± 0.11 | 0.14 | 0.17 | -0.36 ± 0.33 | 0.11 | 0.3100 | **-0.37 ± 0.08** | **0.72** | **0.0018** |
| 3 | 0.37 ± 0.27 | 0.16 | 0.2138 | -0.22 ± 0.13 | 0.20 | 0.1100 | **-0.34 ± 0.09** | **0.79** | **0.0060** |
| 4 | 0.33 ± 0.44 | 0.04 | 0.47 | -0.11 ± 0.20 | 0.02 | 0.5900 | **-0.52 ± 0.08** | **0.84** | **0.0002** |
| 5 | 0.55 ± 0.25 | 0.33 | 0.053 | <u>-0.03 ± 0.14</u> | 0.00 | 0.8000 | **-0.39 ± 0.09** | **0.66** | **0.0023** |
| 6 | <u>0.02 ± 0.27</u> | 0.00 | 0.9427 | **-0.29 ± 0.10** | **0.37** | **0.0120** | **-0.39 ± 0.08** | **0.64** | **0.0003** |
| Fit Average | 0.20 ± 0.11 | 0.21 | 0.11 | **-0.19 ± 0.04** | **0.58** | **0.0005** | **-0.47 ± 0.04** | **0.94** | **<0.0001** |
| Slope Dist ≠ 0 | 0.13 ± 0.14 | - | 0.3789 | **-0.21 ± 0.05** | - | **0.0050** | **-0.45 ± 0.05** | - | **0.0050** |

We found laminar differences in both the average density and the RC spatial pattern of active neurons with odor exposure. In L2, the average density of tdTom(+) neurons (in cells/mm^2^) was significantly greater in odor-exposed animals, NEO (243 ± 43) and HCO (199 ± 13), compared to HC animals (113 ± 18, p: 0.013 KW-test, **Figure 5F1**). In L3, average density did not vary with condition (HC: 52 ± 11; HCO: 68 ± 5, and NEO: 76 ± 15, p: 0.149 KW-test, **Figure 5F2**). In contrast, we found RC spatial patterning of neural activity in L3 (**Figure 5D**) but not L2 (**Figure 5C**). The normalized density of tdTom(+) neurons was plotted against RC distance for individual mice (open triangles) and averaged across animals (solid triangles, **Figure 5C,D**). RC patterning was defined as significant non-zero slope values from linear fits of RC densities in individual animals as well as across animals within a group (**Table 2)**. In L2, the distributions of slope values did not significantly differ from zero (HC: 0.39 ± 0.27; HCO: -0.01 ± 0.10; NEO: -0.08 ± 0.07 mm^-1^, p: 0.065 - 0.38, MWU-test, **Figure 5C, Table 2**) or between conditions (p: 0.104 KW-test, **Figure 5E1**). However, in L3, there was significant RC patterning of active neurons that further differed between HC, HCO and NEO conditions (**Figure 5D, E2**). All NEO mice showed a significant decrease in the density of active neurons along the RC axis (filled red triangles, slope distribution ≠ 0: -0.45 ± 0.05 mm^-1^, **p: 0.005, MWU-test, **Figure 5E2, Table 2**). Further, the distribution of slope values was significantly more negative in NEO mice than HCO or HC (★★, p: 0.0046, KW-test, **Figure 5E2**). Consistent with findings from individual NEO animals, the average change in RC density across animals was also significantly negative (red triangles, -0.47 ± 0.04, p<0.000, **Figure 5D3**). In HCO animals, RC patterning was shallower and less reliable in individual mice than NEO animals. The average density across animals decreased significantly with RC distance but the slope was less than half that of NEO animals (gold triangles: -0.19 ± 0.04; p: 0.0005, linear regression, **Figure 5D2**). Further, although the distribution of slopes across mice was significantly non-zero (-0.21 ± 0.12 mm^-1^, **p: 0.005, MWU-test), RC decreases were rarely significant in individual mice (n=2, filled orange triangles, **Figure 5E2, Table 2**). In HC animals, the average density across animals appears to increase with RC distance (slope: 0.24 ± 0.10 mm^-1^; p: 0.037, linear regression, **Figure 5D1**). However, in individual animals, the distribution of slopes was inconsistent-positive (n=3), negative (n=2) and neutral (n=1) (**Table 2**) and did not significantly differ from zero (slope: 0.13 ± 0.14 mm^-1^, **Figure 5E2**). Finally, across conditions, changes tdTom(+) densities do not correlate with changes in the total number of cells along the RC axis. In 6 mice, two from each group (NEO, HCO, HC) we quantified the RC density of all cells (DAPI stain) in L2 and L3 **(Supplemental Figure 2**). Total cells consistently increased along the RC axis in L2 (slope distribution ≠ 0: 0.54 ± 0.21 mm^-1^, p: 0.0051 MWU) but not in L3 (0.08 ± 0.11 mm^-1^, p: 0.94 MWU, n=6). To summarize, we show that odor exposure increases the density of active neurons in L2 but not L3, and significantly changes the RC spatial patterning of neural activity in L3 but not L2. The lack of consistent, significant RC patterning in individual mice in HC and HCO animals, suggests that spatial patterning within APC is not a reliable feature of odor processing in familiar environments. In contrast, exploration of a novel odor environment (NEO) strongly and reliably changes RC spatial patterning in L3 compared to HCO, and HC contexts. This suggests that space may be an avenue to differentially process odor information depending on context.

In the first section of this study, we show that inhibition of dPCs in L3 increases along the rostral-caudal axis on the spatial scale of millimeters. We find that a disinhibitory circuit mediated by SST-cells supports this gradient through rostrally-biased inhibition of interneurons (**Figure 6A**). Could these RC asymmetries in inhibition play a role in the RC patterning of neural activity in L3 during odor exposure? In NEO animals, neural activity in L3 decreases from rostral to caudal APC over a spatial scale of millimeters (**Figure 5D2**) comparable to that of increasing inhibition in dPCs (**Figure 1F**). In **Figure 6B**, we plot the average normalized decrease in active L3 neurons in NEO mice and the average normalized increase in inhibition of dPCs along the RC axis. We find that the spatial scales of neural activity and inhibition are well matched with opposing slopes (NEO: slope -0.47 ± 0.04 mm^-1^, R: 0.94, p<0.0001; Inhibition: slope 0.67 ± 0.06 mm^-1^, R: 0.98, p: 0.0001). Thus, when inhibition is weakest, neural activity is maximal (rostral) and when inhibition is strongest, neural activity is minimal (caudal). These findings suggest that inhibitory circuitry could underlie RC patterning of neural activity in L3 of APC. Further, the recruitment of inhibitory gradients and may depend on the context of odor experience. In contrast, inhibition is weaker in L2 where the spatial profiles of neural activity are approximately uniform in L2 and do not seem to vary with odor context. Altogether, these laminar and RC differences in inhibition and neural activity suggest spatially dependent and independent mechanisms work in parallel during odor processing in APC.

**Figure 6.**
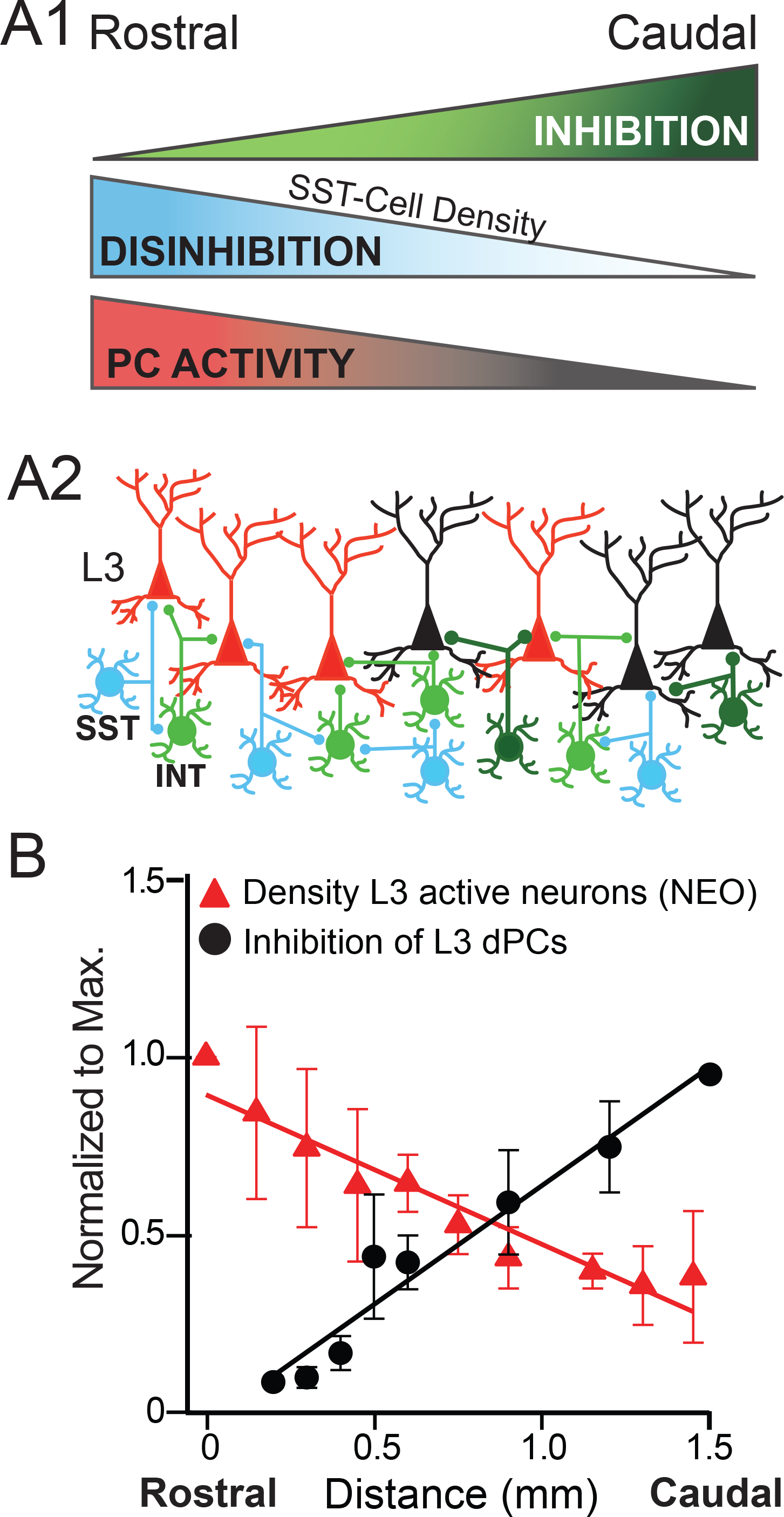
The spatial profiles of inhibition are commensurate with neural activity along the RC axis of APC. **A1)**Schematic summarizing rostral-caudal spatial profiles of inhibition (green), SST-interneuron density and SST-mediated inhibition of interneurons (blue) and active neurons in L3 of NEO mice (red). **A2)** Proposed disinhibitory circuit consisting of a higher density of rostral SST-cells (blue) that inhibit interneurons (light green) and disinhibit PCs increasing rostral neural activity (red PCs). In caudal APC, lower density of SST-cells allows greater inhibition (dark green) and less active PCs (black). **B)** Normalized density of active neurons (red triangles) decreases along the RC axis (slope: -0.47 mm^-1^, R^2^:0.94, p<0.0001, from **Figure 5D3**), as normalized inhibition of dPCs increases (slope: +0.67 mm^-1^, R^2^:0.95, p<0.0001, from **Figure 1F**). Because there are only two points at 1.5 mm in **Figure 1F**, inhibitory strength was normalized to a projected “maximum” strength (8.17 pAs) at 1.6 mm, based on the linear fit to inhibitory strength versus distance.

## Discussion

In this study, we demonstrate rostral-caudal spatial patterning in inhibitory circuitry and neural activity in APC. Our findings reproduce earlier studies that have shown caudally-biased asymmetric inhibition of PCs ^32^ and a RC decline in fos(+) neurons following odor exposure ^22^. However, the underlying circuitry and functional significance of these findings are unknown. Here, we provide three major advances. First, we describe a disinhibitory circuit mediated by SST-cells that decreases rostral inhibition relative to caudal inhibition in L3 PCs. Second, we show that RC patterning of neural activity is confined to L3 and differs with odor exposure in familiar (HCO) versus novel (NEO) contexts. Finally, the density of active neurons decreases along the RC axis following odor exposure in the NEO context commensurate with increasing inhibition of L3 PCs. Specifically, rostral PCs are more active and receive significantly less inhibition (disinhibited) whereas caudal PCs receive stronger inhibition and are less active. Altogether, our findings provide new evidence for RC spatial organization within APC as well as a potential circuit mechanism for varying olfactory processing in different contexts.

### Disinhibition by Somatostatin Interneurons

Inhibition plays a critical role in the processing and representation of sensory information in the cortex ^43^. In APC, inhibition balances excitation ^39,44,45^, narrows synaptic integration windows ^20,44-48^, supports oscillatory activity and sharpens odor tuning ^23,49-51^. Despite the prominent role inhibition plays in shaping cortical responses, few studies have addressed circuits that modulate inhibition in APC ^49,51^. In neocortex, a number of disinhibitory circuits have been implicated in the gating or tuning of cortical responses to sensory stimuli ^52-59^. For example, SST-cells inhibit a range of interneuron classes including fast-spiking, PV interneurons both in neocortex ^56,57^ and APC ^42,51^. In this study, we show that SST interneurons provide rostrally-biased inhibition to L3 interneuron and thus, mediate rostral disinhibition of PCs. Disrupting this disinhibitory circuit through selective optogenetic activation or chemogenetic inhibition of SST cells neutralizes the caudally-biased inhibitory gradient onto dPCs. These findings suggest that SST-cells both inhibit ^42^ and disinhibit PCs in APC and play a role in the RC patterning of inhibition.

Recent studies have shown that SST interneuron activity is modulated in different contexts. In sensory neocortices, SST-cell activity is enhanced by cholinergic modulation ^56,60,61^, running during visual sensory stimulation ^62^, or engagement in an auditory task ^61^. In somatosensory cortex, whisking specifically increases the firing rates of fast-spiking, SST-cells that preferentially inhibit PV cells ^56,59^. Likewise, we have shown that two-thirds of SST cells in APC are FS and strongly inhibit PV-like cells ^42^. Given that sniffing and whisking are correlated ^63,64^, an intriguing possibility is that actively exploring (running, sniffing and whisking) a novel odor environment (NEO) globally enhances SST-cell activity in sensory cortices. We propose that in APC, enhanced SST-cell activity gates rostral disinhibition and increases in rostral neural activity in NEO animals. This interpretation is consistent with recent studies that show interactions between interneurons in network models ^65^ promote context dependent changes in network activity ^58,61^.

### Spatial patterning of neural activity in APC

The spatial patterning is difficult to investigate *in vivo* due to the extent (~1.5 mm) and ventral location of APC. Population imaging shows minimal spatial variation in response to different odors or intensities, but typically only sample L2 neurons across ~300 µm of the RC axis ^25-27^. Multi-site unit recordings broadly sample the RC axis and suggest RC variation in odor-evoked firing rates ^24^ but sample a small proportion of neurons per region. Likewise, intrinsic signal imaging or local field potential (LFP) recordings broadly sample the RC axis and suggest systematic variation along the RC-axis in concentration thresholds ^28^ and oscillatory activity respectively ^13,29,30,66^. But these tools lack the fine resolution to identify the neural circuits contributing to these responses.

To investigate the spatial profiles neural activity *in vivo*, we used TRAP-mice that conditionally express of cre-recombinase linked to the IEG, *c-fos* ^38^. This tool provides sufficient spatial scale to investigate population activity along the entire RC axis at a resolution amenable microcircuit analysis. TRAP-mice are advantageous over traditional IEG methods because cre-recombinase promotes continuous cytoplasmic expression of tdTom independent of initial strength of activation. The limitation is that the temporal window for capturing activity is longer. Labeling is optimal within one hour of 4-OHT injection and declines significantly ~6 hours post injection ^38,67^. Thus, neural densities are expected to be higher in TRAP animals due to enhanced labeling of weak responses and potential spurious labeling over long time windows. To minimize the latter, animals were undisturbed in the dark for 10h following exposure and HC animals provided a baseline for handling and non-specific labeling.

Consistent with previous IEG immunolabeling ^22,68,69^, the density of activated, TRAP-tdTom(+) cells increases significantly in odor-exposed animals (HCO and NEO) compared to HC animals. We find these changes in density are restricted to L2 whereas RC patterning of neural activity occurs in L3. A lack of RC patterning in L2 was initially surprising since L2 sPCs also receive caudally-biased inhibition. However, L2 sPCs receive weaker inhibition than L3 dPCs ^39^ while L2 semilunar cells and interneurons do not receive asymmetric inhibition. This suggests that the uniformly distributed spatial pattern of neural activity in L2 is inherited from the spatial profile of afferent and/or recurrent excitation ^18-20^. In contrast, interneuron densities, particularly SST cells, are greatest in L3 ^40,42^. We show that individual L3 dPCs receive strong inhibition that increases with caudal position along the RC axis with the same spatial scale as decreases in neural activity. These are ideal conditions for inhibition to dictate L3 RC activity patterns. Altogether, laminar differences in inhibition and RC patterning coincide with layer-specific differences PC subclasses and projection targets ^70,71^ and support the premise that parallel processing streams exist in APC.

### Functional roles for RC asymmetries in olfactory processing

Given the seemingly uniform profile of excitation in APC, a surprising finding is that inhibitory strength increases along the rostral-caudal axis. In entorhinal cortex, a dorsal-ventral inhibitory gradient coincides with an increase in PV interneuron density, changes in receptive field size and increased gamma oscillatory power ^72^. We find SST cells rather than PV cells change in density along the RC axis. Since SST-cell inhibition has a subtractive effect on PC odor tuning ^51^, it is possible that SST-mediated inhibition of PCs supports changes in odor tuning across the RC extent of APC. Alternatively, RC patterning of inhibition and neural activity could bias projections from rostral versus caudal APC in different contexts. For example, higher rostral activity during NEO exploration could preferentially increase feedback to the OB ^73^ or output to the OFC ^11,33^. Finally, RC variation in inhibition may also interact with other RC asymmetries to affect the spatial profiles of neural activity in APC. For example, tufted cells afferents are limited to the rostral-ventral APC ^18^ and the overall density of OB afferents decreases along the RC extent of the LOT. Likewise, projections from PPC ^74^ and frontal cortex ^11,34^ also show RC biases. It remains to be determined if the various sources of RC asymmetry work in concert during olfactory processing. Nonetheless, our study adds to a growing body of evidence that, despite the lack of a topographic code for odor identity, space is a relevant dimension in olfactory processing in piriform cortex.

## Methods

### Animals

A number of transgenic mouse lines and crosses were used in this study. VGAT-ChR2 mice (*VGAT-Chr2:* B6.Cg-Tg(Slc32a1-COP4*H134R/EYFP)8Gfng/J) express channelrhodopsin in all interneurons ^75^. The *SST-Cre* (B6:Sst<tm2.1(cre)Zjh>/J) mice were crossed with *Ai32 mice* (B6:129S-Gt(ROSA)26Sortm32(CAG-COP4*H134R/EYFP)Hze/J) or DREADDi mice (B6N.129-*Gt(ROSA)26Sortm1(CAG-CHRM4*,-mCitrine)Ute*/J) to express channelrhodopsin or the inhibitory DREADD, hM4Di ^37^. TRAP mice (FosCre^ERT^: (B6.129(Cg)-Fos(tm1.1(cre/ERT2)Luo/J) were crossed with Ai14 (B6.Cg-*Gt(ROSA)26Sortm14 (CAG-tdTomato)Hze*/J) to conditionally express tdtomato ^38^. All mice are from Jackson Laboratories. Mice were housed in groups of 2-5 animals on a 10:14 light/dark cycle unless otherwise stated. All experiments involved mice of both sexes and age ranges from P20-P300 as indicated.

### Slice preparation

Brain slices of anterior piriform cortex (APC) were prepared from mice aged P19-35. The mice were anesthetized with isoflurane and decapitated. The brain was removed from the skull and immersed in ice cold oxygenated (95% O_2_-5% CO_2_) ACSF (in mM: 125 NaCl, 2.5 KCl, 25 NaHCO_3_, 1.25 NaH_2_PO_4_, 1.0 MgCl_2_, 25 Dextrose, 2.5 CaCl_2_) (all chemicals from Sigma, USA unless otherwise stated). Parasagittal slices (300 μm) were made using a vibratome (Leica Biosystems) in ice cold ACSF. The slices were transferred to warm ACSF (370C) for 30 min and then rested at 20-220C for 1 hour prior to recording (31-350C). All surgical procedures were approved by the University of Pittsburgh IACUC.

### Electrophysiology

Whole cell, voltage and current clamp recordings were performed using a MultiClamp 700B amplifier (Molecular Devices, Union City, CA). Data were low pass filtered (4 kHz) and digitized at 10 kHz using an ITC-18 (Instrutech) controlled by custom software (Recording Artist, https://bitbucket.org/rgerkin/recording-artist) written in IgorPro (Wavemetrics). Recording pipettes (4-10 MΩ) were pulled from borosilicate glass (1.5 mm, outer diameter) on a Flaming/Brown micropipette puller (Sutter Instruments). The series resistance (<20 MΩ) was not corrected. The intracellular solution consisted of (in mM) 130 K-gluconate, 5 KCl, 2 MgCl_2_, 4 ATP-Mg, 0.3 GTP, 10 HEPES, and 10 phosphocreatine, 0.05% biocytin, 4.5 µM QX-314. Recordings were obtained from L2/3 pyramidal cells (PCs), L2 semilunar (SL) cells as well as interneurons in L2/3. Neurons were visualized using infrared-differential interference contrast microscopy (IR-DIC, Olympus). In transgenic mice, interneurons were targeted using fluorescence (YFP) and PCs as the absence of fluorescence. For all neurons, intrinsic subthreshold properties such as input resistance, and time constant were assessed using a series of hyperpolarizing and depolarizing current steps (-50 pA to 50 pA, 1 s duration). Neural identity was confirmed post hoc using intrinsic properties and anatomical analysis of biocytin fills.

### Light stimulation

Blue light (λ=460-488 nm, GFP block, Olympus) for optical stimulation was provided by metal halide lamp (200W, Prior Scientific) passed through the microscope objective (60x, immersion, Olympus). Light pulses were controlled using a mechanical shutter (Sutter Instruments). The light spot was restricted to a ~70 µm diameter (0.5 mW) using the minimum aperture. To obtain the spatial profile of inhibition, interneurons were focally activated in a 5x4 grid pattern while IPSCs were recorded in interneurons or PCs. The horizontal axis of the grid was centered on the recorded neuron with stimulation sites ranging from -300 µm (rostral) to +300 µm (caudal) at 150 µm increments. The vertical axis ranged L1 to L3 in 125 µm increments corresponding to different lamina. Each grid site was stimulated with 2 light pulses (20 ms duration, 100 ms interpulse interval, 15 s between trials). The 20 ms duration was chosen to reliably evoke least one spike and rarely 2 spikes in response to a single pulse of direct somatic stimulation using the 70 µ spot at 0.5 mW ^39^. Grids were repeated 3-7 times per neuron and each grid site was stimulated once every 6 min. Since solely inhibitory neurons are activated and there is little evidence of depolarizing inhibition, polysynaptic responses are unlikely under these recording conditions.

### CNO Application

Stock solutions (10 mM in 0.9% saline) of the DREADD agonist, Clozapine-N-oxide (CNO), were made fresh for each cohort of animals, aliquoted and stored at -200C for up to 2 weeks. On the day of experiment, CNO stock was diluted (20 µM in ACSF) for bath application.

### Analysis of inhibition

Electrophysiology traces of IPSCs are presented as the average across trials (n=3-7) for individual neurons. IPSC strength was taken as the area (pAs) under the first IPSC of the pair of light pulses. The second IPSC was not analyzed due to unreliable AP firing on the second light pulse ^39^. Average PSCs with minimum amplitude of 10 pA were included for analyses; smaller PSCs were not distinguishable from noise and given a value of 0. To compare the spatial profiles of inhibition across animals IPSC amplitudes were normalized to the strength of the maximum IPSC regardless of location in the grid. The rostral-caudal bias was taken as the average normalized inhibition from the caudal sites minus the average inhibition of the rostral sites, divided by the summed inhibition from both sides. The bias metric ranges from -1 (rostral bias) to +1 (caudal bias). Since L1 inhibition was typically weak ^39^ these sites were excluded from the bias metric.

### TRAP-mice

To label active neurons during odor exposure, we used TRAP mice. Briefly, FosCre^ERT^ mice express a tamoxifen-dependent cre-recombinase under the promoter for the activity dependent, immediate early gene *c-fos*. FosCre^ERT^ mice were crossed with Ai14 mice and the offspring conditionally express tdTomato (tdTom) in active neurons upon tamoxifen administration. We used 4-hydroxytamoxifen (4-OHT) (Sigma) because the time window of activation was faster and narrower than tamoxifen ^38^. Doses of 4-OHT were freshly made on the day of injection. Briefly, 4-OHT (15 mg) was dissolved in in 100% ethanol (200µl) by sonication at 37°C (~1 hr). Then peanut oil (1.5 ml, Sigma) was added and ethanol was removed via centrifugation (15 min) and vacuum evaporation (1-2 hrs). The final solution (50mg/kg) was filtered (0.2 µm) and administered by intraperitoneal injection (~150-200 µl per animal) 30 min before odor exposure.

### Odor exposure and behavior

There were three groups of TRAP-mice (P90-300): 1) home cage animals (HC), 2) home cage plus odor (HCO), and 3) novel environment plus odor (NEO). Experiments were done serially, with 3-4 mice per cohort. In each cohort, there was typically at least one mouse per condition. However, there were losses due to death (n=2), poor perfusion (n=2) and insufficient 4-OHT dosage (n=3). Whenever possible, mice from the same litter were used for each cohort or nearly age-matched litters (± 1 week) were used. Mice were singly housed on a 12:12 light/dark cycle and all testing was done ±1 hr from the onset of the dark cycle. Mice were food restricted (90% body weight) as well as handled and weighed daily for 3 days prior to 4-OHT injection and odor exposure. Odor stimuli were isoamyl acetate (Sigma) or ethyl butyrate (Sigma) at 1:100 dilution in mineral oil. For HCO animals, 100 µl of odor was applied to a cotton ball in an open tube that was placed in the cage for 30 min. For NEO mice, filter paper (0.5 x 0.5 cm) was saturated with odor and then buried in a paper cup filled with clean bedding. To encourage exploration, the cup of odorized bedding was placed at the end of one arm of a divided arena (20 x 10 inches) and a blank cup of bedding in the other arm (**Figure 5B2)**. Following 4-OHT administration and exposure, animals were returned to their home cages and undisturbed in the dark for 10-12 hours. Mice were sacrificed 5 days post 4-OHT and neural activity in each context was quantified as the density of tagged, tdTom(+) neurons in L2 and L3 of APC. The average densities of tdTom cells across L2/3 ranged from ~70-400 cells/mm^2^ depending on condition. A minimum average density of 30 cells/mm^2^ was set as a lower threshold for inclusion of an animal in the data set. In three excluded animals, densities were <10 cells/mm^2^ suggesting that 4-OHT dosage was insufficient.

### Anatomy

Mice were given an overdose of ketamine-xylazine. Mice were then perfused transcardially (20 ml/min) with 0.1 M sodium phosphate buffer (PB), followed by 200 ml of 4% paraformaldehyde (PFA) in 0.2M PB. Brains were removed and fixed in 4% PFA overnight at 4°C, then transferred to a sucrose solution. Coronal slices (50 µm) were cut using a freezing microtome maintained in phosphate buffer prior to immunochemistry (anti-PV or anti-CB staining) and/or mounting. Parvalbumin (PV) cells were immunostained using rabbit anti-parvalbumin (PV27, Swant, 1:1000). Calbindin cells were immunostained using rabbit anti-calbindin D-28K (CB38, Swant, 1:1000). In both cases, the secondary was donkey anti-rabbit Alexa-fluor-488 (#A21206 Life Technologies, 1:500). Every other section was mounted using fluoromount to protect fluorescence and minimize background. Sections were imaged on a Nikon Eclipse-Ci microscope at 4x magnification. Illumination was provided by a mercury lamp (Nikon Intensilight) and delivered through appropriate filter blocks for GFP (495 nm) and tdTomato (585 nm). Light intensity and exposure duration (100-400 ms) were optimized for the first section in a series using automated software (Nikon Elements), then maintained for ensuing sections. Sections were photographed using a CCD HD color camera (Nikon DsFi2).

### Cell counts

Neural densities were quantified as number of cells per mm^2^ in laminar regions of interest (ROI) located directly under the lateral olfactory tract in APC. Counts were made in a single focus plane (4x magnification) for each section chosen to maximize the number of cells in focus. Automated counts of somas were obtained based on fluorescence intensity and circularity using Elements Software (Nikon). Two researchers independently verified all counts with at least one blind to condition. In the event of discrepancy, a third individual, blind to condition, performed counts. Every other coronal section (8-15 sections per animal) was analyzed spanning 1-1.5 mm along the rostral-caudal extent of the APC. The average density was taken across all sections in a given animal. To assess rostral-caudal spatial patterning, densities in each section were normalized to the most rostral section corresponding to ~2.46 mm from Bregma ^41^. For each animal, the slope of the least-squares linear regression between normalized density and RC distance was used to quantify spatial patterning.

### Statistics

All data is presented as mean ± SE unless otherwise stated. Initial sample sizes were determined based on previous studies ^22, 57^ using comparable techniques and statistical comparisons. Power analyses were conducted following statistical analysis to ensure sufficient power (Supplemental Tables 1,2,3). Statistical tests were performed using two tailed, one or two-sample, paired or unpaired Student’s t-test as appropriate. In cases of small sample sizes (<10) non-parametric tests were used, including the Mann-Whitney U-test (MWU) for unpaired data and the Wilcoxon Signed Ranks test (WSR) for paired data. For multiple comparisons we used ANOVA with post hoc Tukey Test (ANOVA-Tukey). For groups with small sample sizes multiple comparisons were made using non-parametric, Kruskal-Wallis tests (KW-test). All statistical tests are indicated in the main text and/or figure legends.

## Acknowledgements

We thank Brent Doiron and Nathan Urban for helpful comments on this study and manuscript. We thank Samantha Mielo for technical support. This work was supported by a National Institute on Deafness and Other Communication Disorders Grant (R01 DC015139) to AMMO and an RK Mellon Fellowship to AML.

## Author Contributions

AML and AMMO designed the project, collected and analyzed electrophysiological and anatomical data, and wrote the manuscript. NWV collected and analyzed electrophysiological data. MCB, and PS collected and analyzed anatomical data.

---

**Supplemental Figure 1:**
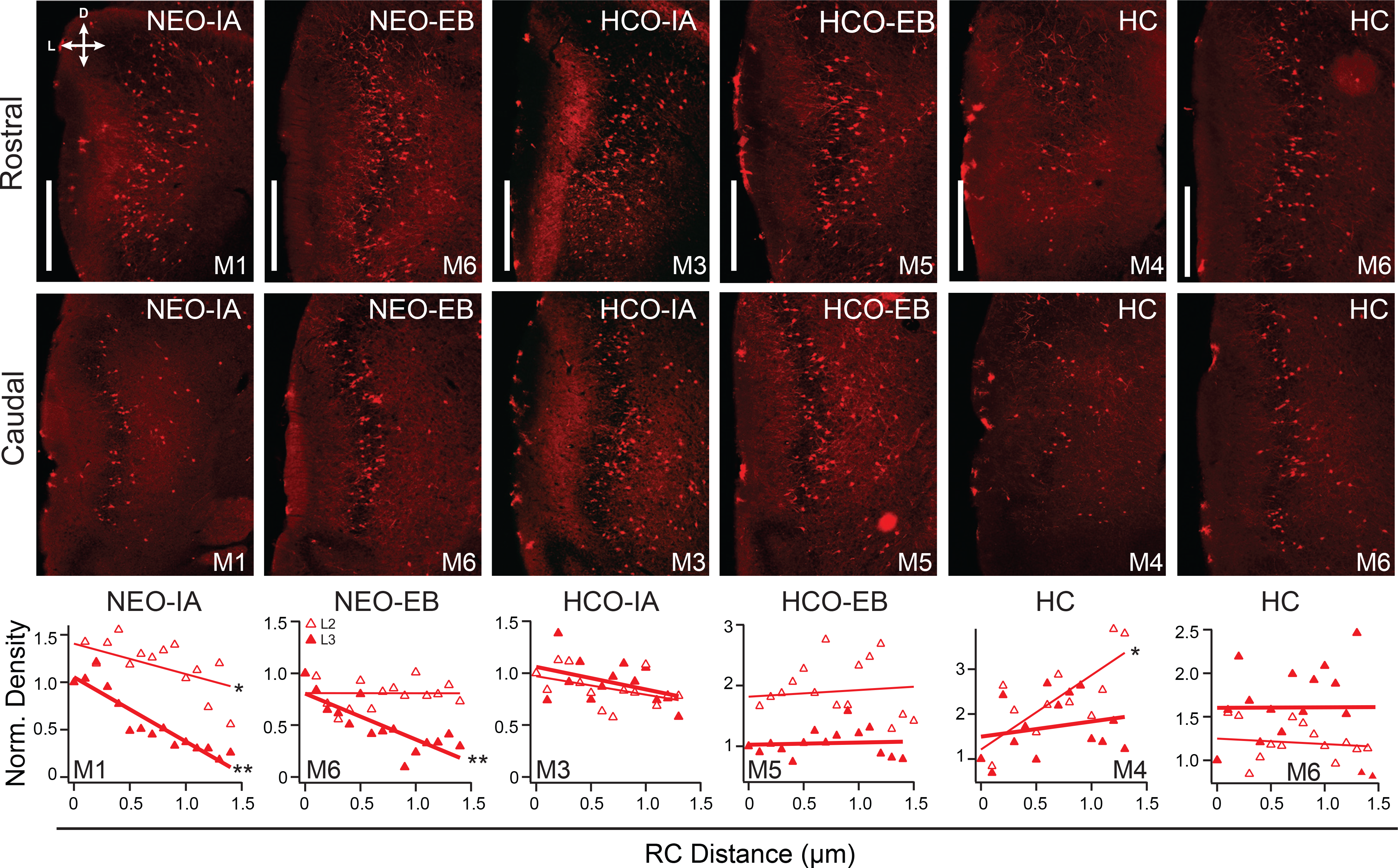
Fos-tdTom(+) cells in anterior piriform cortex (APC). Representative coronal sections from the rostral (0-200 µm, upper row) and caudal (within last 300 µm, middle row) APC showing fos-tdTom(+) cells activated during novel odor exposure in a novel environment (NEO) or the home cage (HCO), or the home cage with no odor (HC). Bottom row: Normalized density (Layer 2, open triangles, L3 filled triangles) versus distance along the RC axis of Fos-tdTom(+) cells for the mouse corresponding to the sections above. Linear regression fits (thin lines, L2: thick lines, L3) with slopes that significantly differed from zero are indicated by astrisks: * p<0.05, or ** p<0.01. Each column is data from an individual mouse and the mouse number (M1-6) matches the mouse numbers in Table 2 (main text). Responses to isoamyl-acetate (IA, left) or ethyl butyrate (EB, right) are shown for each odor condition.

**Supplemental Figure 2:**
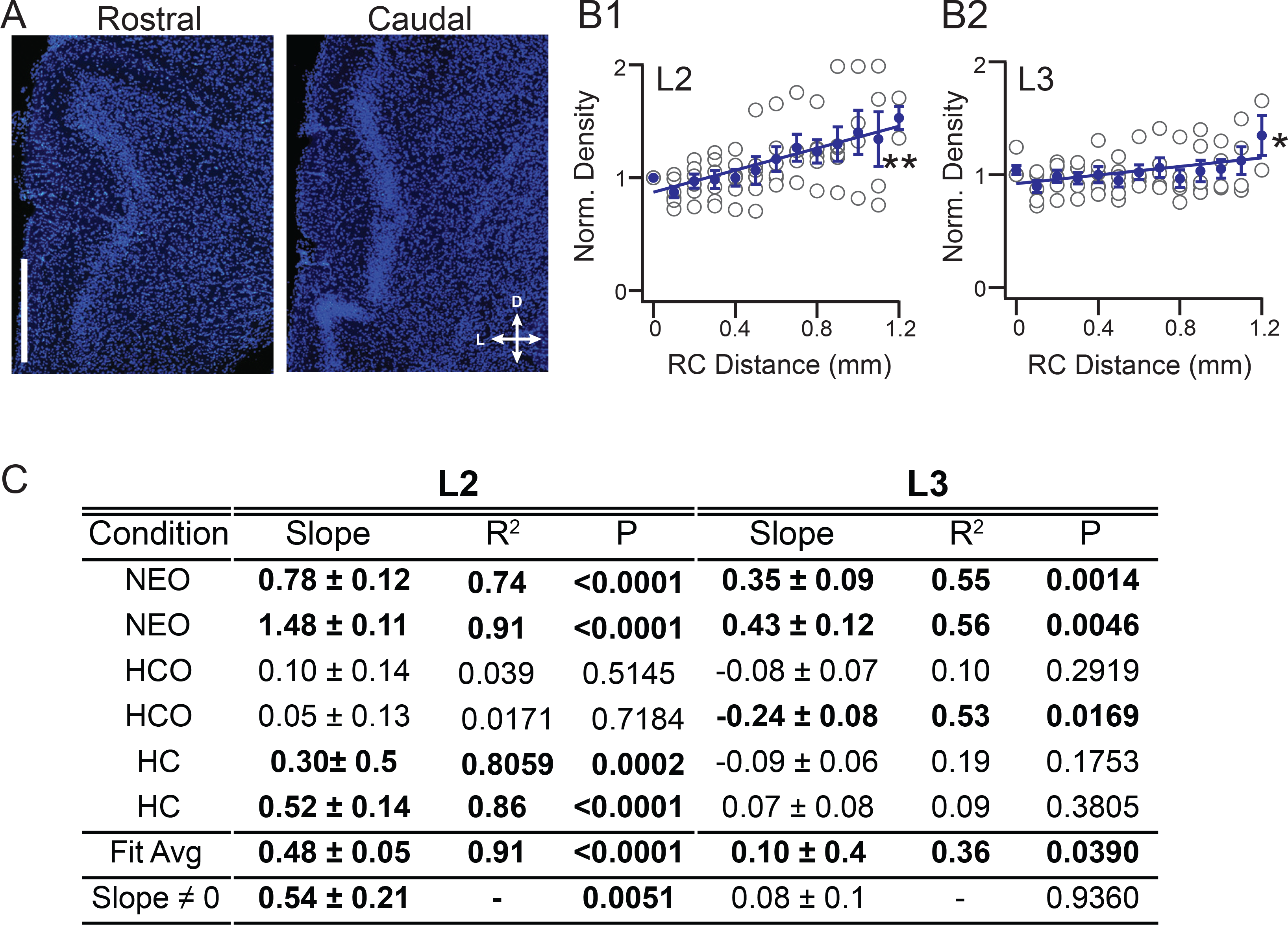
DAPI staining in anterior piriform cortex (APC). Representative coronal sections from rostral (0-200 µm, upper row) and caudal (within last 300 µm) APC showing DAPI(+) nuclei from mice exposed to novel odor exposure in a novel environment (NEO). B) Bottom row: Normalized density versus distance along the RC axis of DAPI(+) cells for individual mice (open circles) and average across mice (blue filled circles) in L2 (B1) and L3 (B2). Linear regression fits of average data have slopes that significantly differed from zero are indicated by astrisks: * p<0.05, or ** p<0.01. C) Table with slope (mm^-1^) data and statistics for each animal. Bold numbers correspond to significant slopes (p<0.05). Fit average corresponds to the linear fits shown in (B). The slope distribution was significantly different from zero for L2 but not L3. HCO: home cage + odor, or the home cage with no odor (HC). Scale: 500 µm

**Supplemental Table 1:** Summary stats for one sample t-test and Mann-Whitney U-test (MWU). M-mean, SD-standard deviation, SE-standard error, n-number of samples, “1-β” power analysis results at α: 0.05 given sample number. Bold: Significant p-values, red : 1-β < 0.7.

| Location | | Distribution $\neq 0$ | mean | SD | Test | p | n | $\alpha: 0.05$<br>$1-\beta$ |
| --- | --- | --- | --- | --- | --- | --- | --- | --- |
| Figure 1 | | bias SL $\neq 0$ | 0.08 | 0.40 | 1 sample t-test | 0.490 | 13 | 0.12 |
| | | bias sPC $\neq 0$ | 0.22 | 0.30 | 1 sample t-test | **0.016** | 14 | 0.85 |
| | | bias dPC $\neq 0$ | 0.19 | 0.20 | 1 sample t-test | **0.001** | 16 | 0.9673 |
| Figure 2 | | bias L2 INT $\neq 0$ | 0.04 | 0.30 | 1 sample t-test | 0.110 | 18 | 0.09 |
| | | bias L3 INT $\neq 0$ | -0.13 | 0.25 | 1 sample t-test | **0.013** | 25 | 0.74 |
| Figure 4 | | bias L3 INT $\neq 0$<br>SST | -0.11 | 0.19 | 1 sample t-test | **0.020** | 22 | 0.81 |
| | | bias dPC $\neq 0$ SST | 0.06 | 0.30 | 1 sample t-test | 0.450 | 14 | |
| | | bias dPC $\neq 0$ CNO | 0.17 | 0.28 | 1 sample t-test | 0.051 | 12 | 0.74 |
| | | bias dPC $\neq 0$ PRE | 0.19 | 0.21 | 1 sample t-test | **0.010** | 12 | 0.91 |
| Figure 3 | TABLE 1 | Slope SST $\neq 0$ | -0.2 | 0.11 | MWU-test | **0.002** | 7 | 0.99 |
| | | Slope PV $\neq 0$ | -0.16 | 0.20 | MWU-test | 0.066 | 6 | 0.54 |
| | | Slope CB $\neq 0$ | 0.13 | 0.39 | MWU-test | 0.370 | 6 | 0.12 |
| Figure 5 | TABLE 2 | Slope HC L2 $\neq 0$ | 0.39 | 0.66 | MWU-test | 0.379 | 6 | 0.3 |
| | | Slope HC L3 $\neq 0$ | 0.13 | 0.34 | MWU-test | 0.379 | 6 | 0.16 |
| | | Slope HCO L2 $\neq 0$ | 0.01 | 0.24 | MWU-test | 0.379 | 6 | 0.05 |
| | | Slope HCO L3 $\neq 0$ | -0.21 | 0.12 | MWU-test | **0.005** | 6 | 0.99 |
| | | Slope NEO L2 $\neq 0$ | -0.08 | 0.17 | MWU-test | 0.065 | 6 | 0.21 |
| | | Slope NEO L3 $\neq 0$ | -0.45 | 0.12 | MWU-test | **0.005** | 6 | 1 |

**Supplemental Table 2:** Summary stats for F-test for non-zero slope of linear regression. n-number of samples, “1-β” power analysis results at α: 0.05 given sample number. Bold: Significant p-values, red : 1-β < 0.7. For density measures, power was analyzed for the mice yielding significant results but the lowest R-values and the lowest number of samples. These represent the minimum power for significant findings within the group.

| Location | | Regression | slope | r | Test | P | n | $\alpha$: 0.05<br>1-β |
| --- | --- | --- | --- | --- | --- | --- | --- | --- |
| Figure 1 |  | Diff IPSC vs Diff Dist | 5.8 | 0.57 | F-test | **0.013** | 19 | 0.7 |
|  |  | IPSC vs RC Dist | 5.4 | 0.64 | F-test | **0.003** | 27 | 1 |
| Figure 3 | Table 1 | SST-M1 | -0.13 | 0.69 | F-test | **0.001** | 18 | 0.9 |
|  |  | SST -M6 | -0.38 | 0.93 | F-test | **0.000** | 12 | 0.9 |
|  |  | SST -AVG | -0.25 | 0.94 | F-test | **0.000** | 10 | 0.9 |
|  |  | PV-M5 | -0.44 | 0.75 | F-test | **0.012** | 10 | 0.75 |
|  |  | PV-Avg | -0.27 | 0.68 | F-test | **0.030** | 10 | 0.6 |
|  |  | CB-M3 | 0.67 | 0.68 | F-test | **0.010** | 13 | 0.75 |
|  |  | CB-M5 | 0.44 | 0.59 | F-test | **0.021** | 15 | 0.67 |
| Figure 5 | Table 2 | HC L2-M4 | 1.62 | 0.78 | F-test | **0.001** | 14 | 0.9 |
|  |  | HC-L2 Avg | 0.42 | 0.76 | F-test | **0.016** | 13 | 0.85 |
|  |  | HC L3 | -0.34 | 0.65 | F-test | **0.022** | 12 | 0.65 |
|  |  | HCO-L2- M4 | 0.3 | 0.52 | F-test | **0.037** | 16 | 0.55 |
|  |  | HCO-L3-M1 | -0.24 | 0.57 | F-test | **0.033** | 14 | 0.55 |
|  |  | HCO-L3-M6 | -0.29 | 0.61 | F-test | **0.012** | 16 | 0.7 |
|  |  | HCO-L3-Avg | -0.19 | 0.76 | F-test | **0.001** | 16 | 0.9 |
|  |  | NEO-L2-M1 | -0.32 | 0.54 | F-test | **0.028** | 15 | 0.55 |
|  |  | NEO-L2-M5 | -0.1 | 0.79 | F-test | **0.003** | 11 | 0.85 |
|  |  | NEO-L3-M3 | -0.34 | 0.79 | F-test | **0.006** | 10 | 0.8 |
| Figure 5, 6, Table 2 |  | NEO-L3-Avg | -0.47 | 0.97 | F-test | **0.000** | 14 | 0.95 |
| Figure 6 |  | IPSC vs RC Dist | 0.67 | 0.97 | F-test | **0.000** | 27 | 1 |

**Supplemental Table 3:**
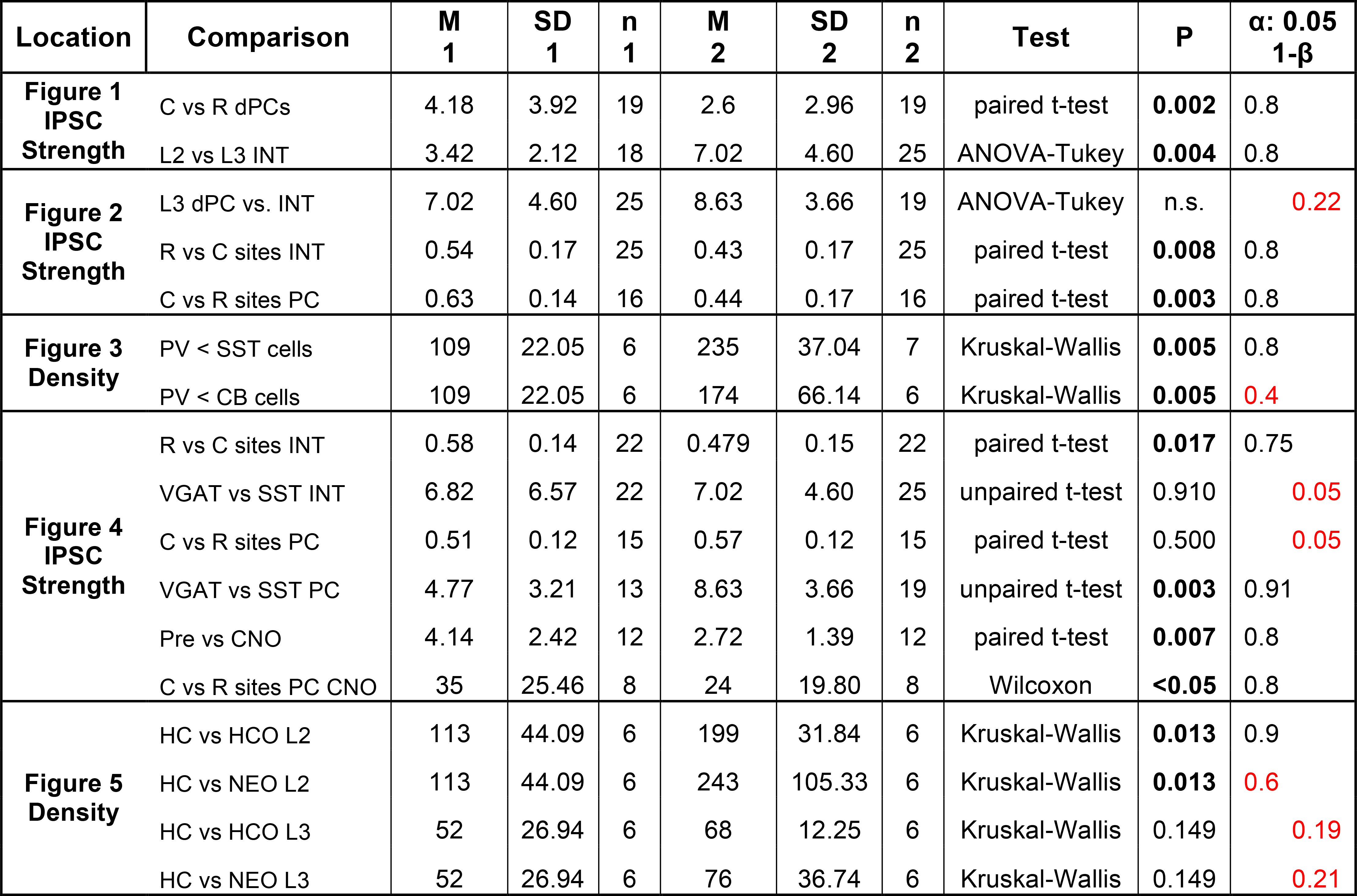
Summary stats for two sample comparisons. M-mean, SD-standard deviation, n-
number of samples, “1-β” power analysis results at α: 0.05 given sample number. Bold: Significant p-values, red : 1-β < 0.7. Abbreviations: C: Caudal, R: Rostral, INT: interneurons, PC: pyramidal cell, PV: parvalbumin, SST: somatostatin, CB: Calbindin, CNO: clozapine-n-oxide, HC: home cage no odor, HCO: Home cage plus odor, NEO: novel environment plus odor, L2: layer 2, L3: Layer 3. M: Mean, n: number of samples.

| Location | Comparison | M<br>1 | SD<br>1 | n<br>1 | M<br>2 | SD<br>2 | n<br>2 | Test | P | $\alpha$: 0.05<br>1-β |
| --- | --- | --- | --- | --- | --- | --- | --- | --- | --- | --- |
| **Figure 1**<br>**IPSC**<br>**Strength** | C vs R dPCs | 4.18 | 3.92 | 19 | 2.6 | 2.96 | 19 | paired t-test | **0.002** | 0.8 |
|  | L2 vs L3 INT | 3.42 | 2.12 | 18 | 7.02 | 4.60 | 25 | ANOVA-Tukey | **0.004** | 0.8 |
| **Figure 2**<br>**IPSC**<br>**Strength** | L3 dPC vs. INT | 7.02 | 4.60 | 25 | 8.63 | 3.66 | 19 | ANOVA-Tukey | n.s. | 0.22 |
|  | R vs C sites INT | 0.54 | 0.17 | 25 | 0.43 | 0.17 | 25 | paired t-test | **0.008** | 0.8 |
|  | C vs R sites PC | 0.63 | 0.14 | 16 | 0.44 | 0.17 | 16 | paired t-test | **0.003** | 0.8 |
| **Figure 3**<br>**Density** | PV < SST cells | 109 | 22.05 | 6 | 235 | 37.04 | 7 | Kruskal-Wallis | **0.005** | 0.8 |
|  | PV < CB cells | 109 | 22.05 | 6 | 174 | 66.14 | 6 | Kruskal-Wallis | **0.005** | 0.4 |
| **Figure 4**<br>**IPSC**<br>**Strength** | R vs C sites INT | 0.58 | 0.14 | 22 | 0.479 | 0.15 | 22 | paired t-test | **0.017** | 0.75 |
|  | VGAT vs SST INT | 6.82 | 6.57 | 22 | 7.02 | 4.60 | 25 | unpaired t-test | 0.910 | 0.05 |
|  | C vs R sites PC | 0.51 | 0.12 | 15 | 0.57 | 0.12 | 15 | paired t-test | 0.500 | 0.05 |
|  | VGAT vs SST PC | 4.77 | 3.21 | 13 | 8.63 | 3.66 | 19 | unpaired t-test | **0.003** | 0.91 |
|  | Pre vs CNO | 4.14 | 2.42 | 12 | 2.72 | 1.39 | 12 | paired t-test | **0.007** | 0.8 |
|  | C vs R sites PC CNO | 35 | 25.46 | 8 | 24 | 19.80 | 8 | Wilcoxon | **<0.05** | 0.8 |
| **Figure 5**<br>**Density** | HC vs HCO L2 | 113 | 44.09 | 6 | 199 | 31.84 | 6 | Kruskal-Wallis | **0.013** | 0.9 |
|  | HC vs NEO L2 | 113 | 44.09 | 6 | 243 | 105.33 | 6 | Kruskal-Wallis | **0.013** | 0.6 |
|  | HC vs HCO L3 | 52 | 26.94 | 6 | 68 | 12.25 | 6 | Kruskal-Wallis | 0.149 | 0.19 |
|  | HC vs NEO L3 | 52 | 26.94 | 6 | 76 | 36.74 | 6 | Kruskal-Wallis | 0.149 | 0.21 |

